# Deep Bayesian networks for uncertainty estimation and adversarial resistance of white matter hyperintensity segmentation

**DOI:** 10.1101/2021.08.18.456666

**Authors:** Parisa Mojiri Forooshani, Mahdi Biparva, Emmanuel E. Ntiri, Joel Ramirez, Lyndon Boone, Melissa F. Holmes, Sabrina Adamo, Fuqiang Gao, Miracle Ozzoude, Christopher J.M. Scott, Dar Dowlatshahi, Jane M. Lawrence-Dewar, Donna Kwan, Anthony E. Lang, Karine Marcotte, Carol Leonard, Elizabeth Rochon, Chris Heyn, Robert Bartha, Stephen Strother, Jean-Claude Tardif, Sean Symons, Mario Masellis, Richard H. Swartz, Alan Moody, Sandra E. Black, Maged Goubran

## Abstract

White matter hyperintensities (WMH) are frequently observed on structural neuroimaging of elderly populations and are associated with cognitive decline and increased risk of dementia. Many existing WMH segmentation algorithms produce suboptimal results in populations with vascular lesions or brain atrophy, or require parameter tuning and are computationally expensive. Additionally, most algorithms do not generate a confidence estimate of segmentation quality, limiting their interpretation. MRI-based segmentation methods are often sensitive to acquisition protocols, scanners, noise-level, and image contrast, failing to generalize to other populations and out-of-distribution datasets. Given these concerns, we propose a novel Bayesian 3D Convolutional Neural Network (CNN) with a U-Net architecture that automatically segments WMH, provides uncertainty estimates of the segmentation output for quality control and is robust to changes in acquisition protocols. We also provide a second model to differentiate deep and periventricular WMH. 432 subjects were recruited to train the CNNs from four multi-site imaging studies. A separate test set of 158 subjects was used for evaluation, including an unseen multi-site study. We compared our model to two established state-of-the-art techniques (BIANCA and DeepMedic), highlighting its accuracy and efficiency. Our Bayesian 3D U-Net achieved the highest Dice similarity coefficient of 0.89 ± 0.08 and the lowest modified Hausdorff distance of 2.98 ± 4.40 mm. We further validated our models highlighting their robustness on ‘clinical adversarial cases’ simulating data with low signal-to-noise ratio, low resolution, and different contrast (stemming from MRI sequences with different parameters). Our pipeline and models are available at: https://hypermapp3r.readthedocs.io

## Introduction

### Clinical Motivation

White matter hyperintensities (WMH) are commonly observed MRI-based biomarkers of cerebral small vessel disease and have been associated with aging and neurodegenerative diseases such as Parkinson’s disease (PD) and Alzheimer’s disease (AD) (S. Lee et al. 2016; Prins and Scheltens 2015; S.-J. Lee et al. 2010). WMH have been linked to global cognitive impairment (Jokinen et al. 2020), a decline in mental processing speed (van den Heuvel 2006), increased risk of late onset depression (Herrmann, Le Masurier, and Ebmeier 2008), and increased risk of stroke, dementia, and mortality (Debette and Markus 2010). Since WMH often occurs in the preclinical stage of dementia, it may represent one of the pathological processes indicating progression from mild cognitive impairment (MCI) to dementia (Smith et al. 2008) and should be considered as a covariant of interest at baseline and longitudinally in future AD treatments (Carmichael et al. 2010). Furthermore, periventricular venous collagenosis is associated with WMH in both AD and non-AD patients (Keith et al. 2017).

WMH appear as hyperintense (bright) on both T2-weighted (T2w) and fluid attenuated inversion recovery (FLAIR) MRI images and hypointense (dark) on T1-weighted (T1w) images. While T1w-based WMH estimates have been shown to have some degree of correlation with estimates based on T2w and FLAIR image (Dadar et al. 2018), the inclusion of the two sequences provides improved contrast and visualization of WMH borders, and thus a more accurate segmentation and volume quantification. This is especially important for the classification of both deep WMH (dWMH, found in deep white matter) and periventricular WMH (pvWMH, which extend from the ventricular wall). dWMH presence has been linked to the incidence of migraine (Hong et al. 2020), while increases in pvWMH have been associated with AD progression (Kilgore et al. 2020), as well as stroke (Hernández, Piper, and Bastin 2014). There has been debate in the literature regarding whether or not WMH should be differentiated based on their location, or if they should be considered to be the same based on continuous spectrum (Wardlaw, Valdés Hernández, and Muñoz-Maniega 2015). The exploration of WMH involvement in neurodegenerative disease warrants an accurate segmentation, quantification and classification of WMH and its subtypes.

### Related Works

#### Automated segmentation

There has been a large amount of work revolving around WMH segmentation in clinical and neuroscience research. Traditionally, manual tracings are employed as the gold standard. Individuals trained to segment WMH are aware of the features that distinguish WMH from normal-appearing white matter and other tissue types. The large amount of time and training required to manually segment WMH necessitate the need for automatic methods to segment WMH. Despite the bevy of available segmentation methods (Griffanti et al. 2016; Zhang et al. 2020; Y. Wang et al. 2012), several challenges still exist, such as the lack of uniformity across different image types and differences in imaging protocols/scanners, as well as the lack of methods for quality control. For example, segmentation methods that rely on image intensity (Yoo et al. 2014; Griffanti et al. 2016) had varying degrees of success. While methods that focus on thresholding based on intensity often fail to detect smaller, lower intensity dWMH, thereby increasing the rate of false positives (Jeon et al. 2011). In addition, datasets used for training and testing are often small, and may not always be tested across different scanner types.

Supervised methods utilize previously acquired ground truth, capitalizing on features from the data to produce accurate segmentations on other datasets. Many state-of-the-art (SOTA) WMH segmentation methods are based on a supervised learning formulation. BIANCA, short for “Brain intensity abnormality classification algorithm” (Griffanti et al. 2016) is a fully automated supervised method based on the k-nearest neighbor (kNN) algorithm. The LOCally Adaptive Threshold Estimation (LOCATE) method was later proposed (Sundaresan et al. 2019) to determine local thresholds that are sensitive to spatial differences in the lesion probabilities to improve BIANCA’s binary lesion masks.

In recent years, deep learning with convolutional neural networks (CNNs) has achieved SOTA performance for many medical image segmentation tasks, including WMH and stroke lesions (Guerrero et al. 2018). Kamnitsas et al. (Kamnitsas et al. 2017) proposed a network architecture consisting of two parallel convolutional pathways that processes the 3D input patches at multiple scales, followed by post-processing using a 3D densely connected conditional random field (CRF) to remove false positives. Their method was originally proposed for ischemic stroke and tumor segmentation, but it can be adopted for different lesion segmentations. The U-Net model architecture (Ronneberger, Fischer, and Brox 2015) has been widely used in segmenting biomedical images due to its performance and efficiency of using GPU memory. A CNN model for WMH segmentation that distinguishes between WMH and stroke was presented by Guerrero et al. (Guerrero et al. 2018). Li et al. (Li et al. 2018) presented a U-Net and an ensemble of models trained with random weight initializations to reduce overfitting and boost segmentation results. A skip connection U-Net model was proposed by Wu et al. (Wu et al. 2019) to capture more features and improve the model’s receptive field and was evaluated on WMH segmentation.

#### Uncertainty and Bayesian networks

Uncertainty estimation is critical for understanding the reliability of segmentation networks, and for providing a quantitative assessment of confidence in their outputs. This is specifically important for medical imaging in a clinical setting. Current approaches do not provide uncertainty estimates for their segmentation results. Several studies have investigated uncertainty estimation for deep neural networks (Kendall and Gal 2017; Kendall, Badrinarayanan, and Cipolla 2017; G. Wang et al. 2019). As suggested by Kendall and Gal, there are two major types of predictive uncertainties for deep CNNs: epistemic (model) uncertainty and aleatoric (image-based) uncertainty. Epistemic uncertainty describes limitations in the learning procedure due to limited training data. Aleatoric uncertainty depends on noise or randomness in the input image.

Bayesian Neural Networks (BNNs) have been used to estimate model uncertainty; however, they are hard to implement and are computationally expensive. Previous works have used Stochastic Variational Gradient Descent (SVGD) to perform approximate Bayesian inference on uncertain CNN parameters (Zhu and Zabaras 2018). Other approximation methods have been developed such as Markov Chain Monte Carlo (Neal 2012) that are not scalable for large neural networks with millions of parameters, and variational methods (Graves 2011) to provide an analytical approximation to the posterior probability of unobserved variables, to perform statistical inference over these variables. Bayes by Backprop (BBB) is another approach that combines variational inference with traditional backpropagation to efficiently find the best approximation to the posterior (Blundell et al. 2015). Other studies have used ensembles of multiple models to generate uncertainty (Lakshminarayanan, Pritzel, and Blundell 2016). Alternatively, Monte Carlo (MC) Dropout has been used to demonstrate that dropout at test time can be cast as approximate Bernoulli variational inference to allow an efficient approximation of the model’s posterior distribution without additional parameters (Kendall and Gal 2017).

#### Generalization and Robustness

Machine learning algorithms are usually evaluated by the model’s generalization and robustness (Paschali et al. 2018). Generalization refers to the model performance on an unseen dataset. To build a deep learning model that generalizes well, a large and diverse amount of data is required to avoid overfitting (LeCun, Bengio, and Hinton 2015). This is a significant obstacle for using deep learning in the medical domain, where producing high-quality labeled data is time-consuming, expensive, and requires expert knowledge. Robustness refers to the ability of a model to correctly classify previously unseen examples with noise and slight perturbations, which are more challenging to classify and segment (Rozsa, Gunther, and Boult 2018). MR-based segmentation methods are mostly sensitive to acquisition protocols, scanners, noise-level and image contrast. Paschali et al. (Paschali et al. 2018) investigated the robustness of a variety of medical imaging networks on adversarial cases and found that models that achieve comparable performance in generalizability may have significant differences in their relative exploration of the underlying data manifold, therefore resulting in varying robustness and model sensitivities. Data augmentation can be used as a regularization strategy to control overfitting and improve the model’s robustness and performance (Nalepa, Marcinkiewicz, and Kawulok 2019). Common data augmentation schemes for segmentation of neuroimaging data include affine transformations, elastic deformation, random cropping, flipping about a spatial axis, adding noise and adjusting image contrast.

### Contributions

In this work, we present several innovative methodological and experimental contributions. First, we developed and evaluated a 3D U-Net with MC Dropout as a Bayesian network to segment WMH and provide uncertainty quantification, generating an estimate of the model’s confidence of the predicted segmentation. Second, a separate model was developed to distinguish and segment dWMH and pvWMH, respectively using initial WMH segmentation. Third, unlike previous work, we employed an unseen multisite study for testing, in addition to the four large multi-site datasets with different diagnostic groups used for training. Fourth, we compared our methods against established SOTA techniques on a wide spectrum of white matter disease burden including very mild WMH cases with small volumes which are difficult to capture and segment. Finally, the network was trained using an augmentation scheme that included permutations to noise level, resolution, and contrast to achieve resistance to common changes due to different MRI acquisition protocols and scanners. This adversarial resistance was validated against simulated challenges in clinical datasets, referred to here as “clinical adversarial attacks”. Our trained models are publicly available, and we developed an easy-to-use pipeline with a graphical user interface for making them accessible to users without programming knowledge.

## Materials and Methods

### Participants

To train the WMH segmentation model, a total of 432 subjects were recruited from four multicenter studies: 160 subjects with cerebrovascular disease +/− vascular cognitive impairment (CVD +/− VCI) or PD (55-86, 75% male) through the Ontario Neurodegenerative Disease Research Initiative (ONDRI) (Farhan et al. 2017), 203 individuals with non-surgical carotid stenosis (47-92, 61% male) through the Canadian Atherosclerosis Imaging Network (CAIN) study (ClinicalTrials.gov: NCT01440296), 37 subjects with nonfluent progressive aphasia, semantic dementia (SD) and normal controls through the Language Impairment in Progressive Aphasia (LIPA) study (Marcotte et al. 2017) (55-80), and 32 subjects with CVD, VCI or Alzheimer’s disease (AD) through the Vascular Brain Health (VBH) study (Swardfager et al. 2017) (46-78, 50% male). Ground truth segmentations for WMH were generated using SABRE-Lesion Explorer semi-automated pipeline that generates intensity-based segmentations which are then manually edited by expert annotators trained by a neuroradiologist with an intraclass correlation of ≥ 0.9 (J. Ramirez et al. 2011; Joel Ramirez et al. 2014)). Sequences used for the generation of the ground truth included 3D T1w and T2w FLAIR to accurately delineate the WMH. Participant demographics, diagnosis, Montreal Cognitive Assessment (MoCA) scores, and volumes for the ground truth segmentations and vascular lesions are summarized in **Table 1**.

**Table 1.** Participant demographics, clinical diagnosis, MoCA scores, ICV and ventricle volumes, WMH volumes, and MRI parameters in the training and test datasets. These datasets are used for training and testing SOTA.

|  |  | Training Dataset<br>n = 432* |  |  |  | Test Dataset<br>n = 158 |  |  |  |  |
| --- | --- | --- | --- | --- | --- | --- | --- | --- | --- | --- |
|  |  | CAIN<br>n=203 | ONDRI<br>n= 160 | LIPA<br>n=37 | VBH<br>n=32 | CAIN<br>n=50 | ONDRI<br>n=37 | MITNEC<br>n=53 | LIPA<br>n=10 | VBH<br>n=8 |
| **Demographics** |  |  |  |  |  |  |  |  |  |  |
| Diagnostic Group |  | CN | PD, CVD | CN, FTD | AD, CVD, CN | CN | PD, CVD | MCI, AD, CVD | CN, FTD | AD, CVD, CN |
| Age (years) |  | 70.4 (7.8) | 68.3 (6.9) | 67.0 (7.3) | 76.4 (9.0) | 69 (7.5) | 67.9 (5.6) | 77 (8.7) | 67.9 (12.1) | 77.5 (11.7) |
| Sex (n, % female) |  | 79 (39.0) | 40 (25.0) | -- | 14 (43.8) | 22 (44.0) | 12 (31.5) | 24 (45.2) | -- | 4 (50.0) |
| MOCA total score |  | -- | 25.9 (2.8) | -- | -- | -- | 25.3 (2.9) | 22.2 (4.8) | -- | -- |
| **Neuroimaging Metrics** |  |  |  |  |  |  |  |  |  |  |
| ICV (cc) |  | 1241.5 (118.5) | 1286.1 (135.8) | 1255.7 (132.1) | 1253.1 (171.0) | 1234.9 (140.0) | 1264.38 (151.0) | 1225.8 (174.2) | 1254.5 (153.9) | 1291.2 (149.6) |
| vCSF (cc) |  | 36.9 (19.7) | 38.8 (19.9) | 44.3 (20.0) | 56.5 (37.2) | 36.8 (19.7) | 33.1 (18.1) | 53.3 (21.6) | 45.1 (18.2) | 48.3 (21.3) |
| pvWMH (cc) |  | 7.7 (9.3) | 7.2 (10.1) | 6.7 (9.9) | 20.1 (22.6) | 6.2 (4.9) | 5.5 (7.9) | 32.5 (22.0) | 2.8 (1.6) | 33.6 (35.2) |
| dWMH (cc) |  | 1.1 (1.6) | 0.7 (1.0) | 0.6 (0.7) | 2.1 (2.4) | 0.8 (1.1) | 0.6 (0.8) | 2.2 (1.2) | 0.5 (0.5) | 2.4 (2.3) |
| **MRI acquisition parameters** |  |  |  |  |  |  |  |  |  |  |
| T1 (SPGR) | In-plane Resolution (mm) | 0.94 x 1.1 | 1 x 1 | 0.9 x 0.9 |  | 0.94 x 1.1 | 1 x 1 |  | 0.9 x 0.9 |  |
|  | Slice thickness (mm) | 1.4 | 1 | 1 |  | 1.4 | 1 |  | 1 |  |
|  | FOV (mm) | 240 | 256 | 220 |  | 240 | 256 |  | 220 |  |
| FLAIR | In-plane Resolution (mm) | 1 x 1.1 | 0.94 x 0.94 | 0.9 x 0.9 |  | 1 x 1.1 | 0.94 x 0.94 |  | 0.9 x 0.9 |  |
|  | Slice thickness (mm) | 3 | 3 | 3 |  | 3 | 3 |  | 3 |  |
|  | FOV (mm) | 230 | 240 | 220 |  | 240 | 240 |  | 220 |  |
| Filed strength |  | 3T | 3T | 3T |  | 3T | 3T |  | 3T |  |

\* 378 used for training and 54 for validation during training
Data is presented as mean ± standard deviation unless otherwise specified. ICV= intracranial volume; MoCA=Montreal Cognitive Assessment; vCSF= ventricular cerebrospinal fluid; pvWMH=periventricular white matter hyperintensities; dWMH=deep white matter hyperintensities; CN=cognitively normal; PD=Parkinson's Disease; VCI=vascular cognitive impairment; MCI=mild cognitive impairment; AD=Alzheimer's disease; CVD=cerebrovascular disease; FTD=Frontotemporal dementia.

The models were tested on a total of 158 subjects, including 53 with severe WMH burden (Fazekas 3/3) from the Medical Imaging Trials NEtwork of Canada (MITNEC) Project C6 (ClinicalTrials.gov: NCT02330510), a separate fifth multicenter (unseen) study not part of those used for training. The breakdown of subjects from the five studies is presented in **Table 1**.

### Data preprocessing

The preprocessing techniques used in this study are described in our prior work (Goubran et al. 2019). Briefly, we conducted the following data preprocessing steps on all images prior to training: 1) bias-field correction for B1 inhomogeneities using the *N4* algorithm (Tustison et al. 2010), 2) skull-stripping to separate brain from non-brain tissues (Ntiri et al. 2021), 3) background-cropping using a bounding box such that all voxels outside the bounding box are zero-valued, and 4) Z-score intensity normalization using the mean as the subtrahend and the standard deviation as the divisor for each patient volume.

### Bayesian 3D CNN architecture

Our networks are based on the U-Net architecture (Ronneberger, Fischer, and Brox 2015; Çiçek et al. 2016), which consists of encoder and decoder pathways, with pooling and upsampling operations. The architecture of the network (**Figure 1A**) was based on the original U-Net (Çiçek et al. 2016; Ronneberger, Fischer, and Brox 2015), with some modifications in our 3D implementation. Similar to our prior work (Goubran et al. 2019), residual blocks were added to each encoding layer. Residual blocks resolve the gradient degradation problem that occurs with deeper networks with an increasing number of layers. Also similar to our previous work (Ntiri et al. 2021), dilated convolutions were used which help to enlarge the field of view (FOV) of convolutional filters without losing resolution or coverage (Yu and Koltun 2015). Instance normalization was used instead of batch normalization (Ioffe and Szegedy 2015) to avoid instability associated with batch normalization due to the stochasticity generated by small batch sizes. The network has a depth of 5 layers and 16 initial filters. Here, motivated by approximate Bayesian formulations in deep learning (Gal and Ghahramani 2016), Monte Carlo (MC) dropout layers were added to the network as one of the main contributions of the proposed network. To turn our baseline CNN into a Bayesian CNN, we added MC dropout layers in residual blocks after the first dilated convolution layer which is equivalent to placing a Bernoulli distribution over the weights.

**Figure 1.**
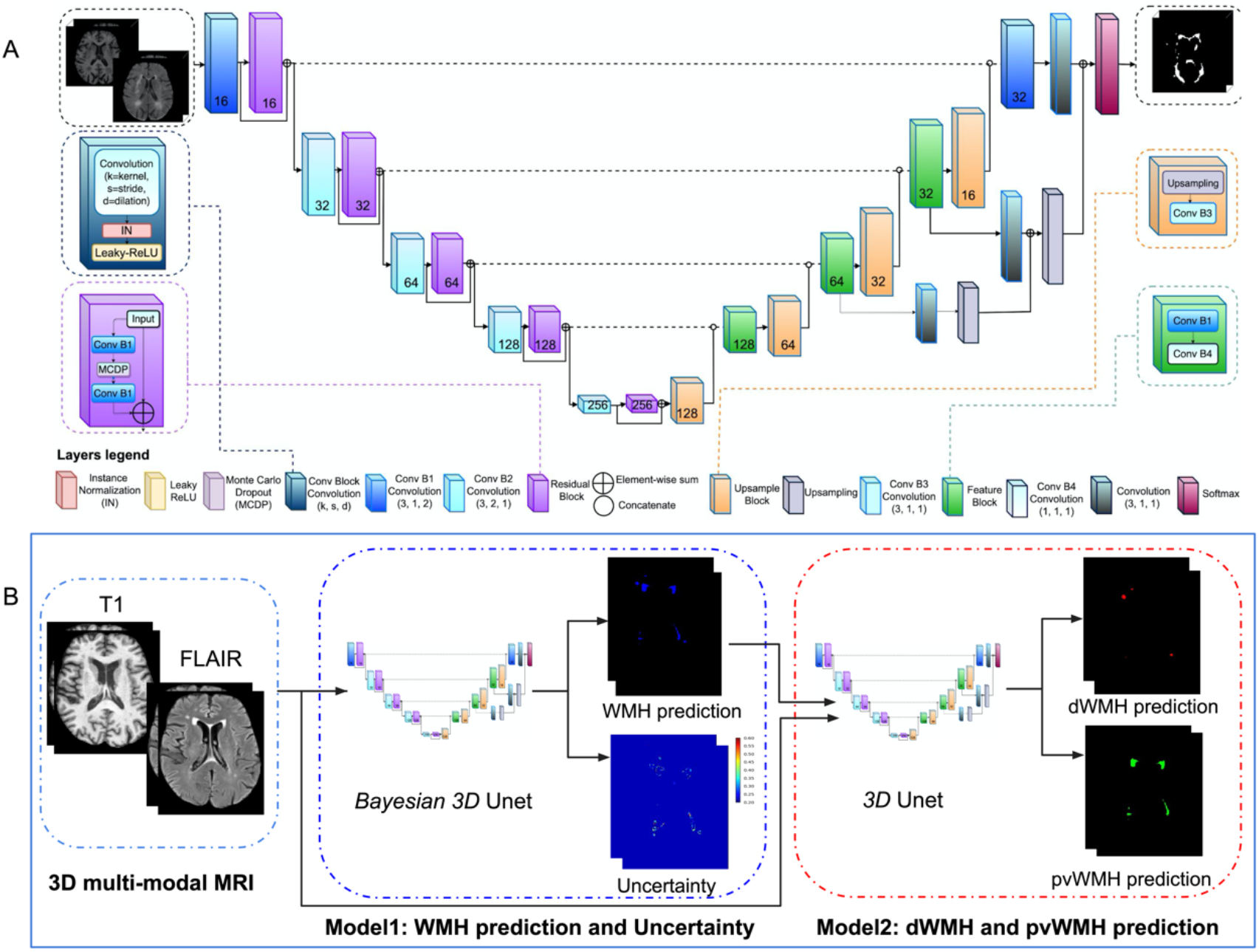
A) Proposed architecture for the Bayesian 3D U-Net convolutional neural network with residual blocks and dilated convolutions. B) Overall inference pipeline to generate WMH segmentation and uncertainty maps as well as a second network to differentiate dWMH and pvWMH.

To optimize the location and rate of MC dropout, validation experiments were performed, testing MC dropout on 1) all layers (both encoder and decoder), 2) encoder layers, 3) three central layers both encoder and decoder, and using dropout rates of 0.3 and 0.5. Based on these experiments, we observed that adding dropout on the encoder layers (the residual blocks) with a dropout rate of 0.3 produced the best results on the validation data.

Each encoder layer consists of a convolutional layer and a residual block (He et al. 2016). At each residual block, the input is split into two paths. The first path consisted of two dilated convolutional layers with kernel sizes 7×7×7 with a MC dropout layer with a rate of 0.3 between the two convolutions, while an identity map was applied to the input data in the second path. Element-wise addition was then applied to the results of the first and second paths, combining the outputs of the two.

At each decoder layer before the final layer, the upsampling block from the previous level is concatenated with the corresponding features on the encoder level. Upsampling modules consisted of an upsampling layer of size 3×3×3, followed by a convolutional layer. The output from the upsampling module was concatenated with the summation output from the respective later on the contracting side, before being passed to a feature block. The feature block consisted of two convolutional layers (one dilated convolution) with a stride of 1×1×1, and kernels of size 7×7×7 and 1×1×1, respectively. The number of filters was halved at each decoder step. At the last layer, the concatenated feature map is passed to a sigmoid function to generate a probability map for the class/voxels of interest.

### Adversarial resistance and augmentation experiments

Data augmentation is an effective way to enlarge the size and quality of training data to improve model generalizability and avoid overfitting due to a limited dataset. Augmentation can equip a deep network with desired invariance and robustness properties. A large list of data augmentation strategies/schemes were evaluated to improve the adversarial resistance of our WMH model to common changes stemming from MRI sequences with different parameters, protocols, and scanners. The effects of three types of augmentations on model performance were evaluated: 1) Geometrical affine transformations, 2) Histogram-based transformations and 3) Pixel-level transformations. Affine transformations included flipping, changes in orientations, random cropping, and random scaling. Histogram-based transformations included histogram equalization, scaling and brightness modification, while pixel-level transformations included the addition of Rician noise (Gudbjartsson and Patz 1995), contrast shifting/scaling and gamma intensity transformations. Based on these experiments an optimal set of augmentations including four transformations per scan were performed on each of the subjects, including flipping along the horizontal axis (Left-Right axis), random rotating by an angle *α* = ±90 along *y* and *z* axis, introducing Rician noise generated by applying the magnitude operation to images with added complex noise, where each channel of the noise is independently sampled from a Gaussian distribution with random standard deviation *σ* = (0.01,0.2), and changing image intensity by gamma *γ* = (0.1,0.5) such that each pixel/voxel intensity is updated as:

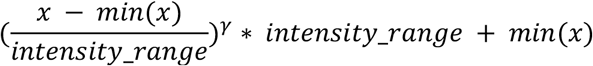

where *intensity_range* is *max*(*x*) – *min*(*x*).

### Loss function

An “equally weighted” formulation of the Dice coefficient was used as the loss function to mitigate the class imbalance issue, in which the majority of voxels in the image do not represent the structure of interest (Milletari, Navab, and Ahmadi 2016). The Dice coefficient is a measure of similarity, determined by a calculation of the overlap between two binary images. Given a predicted binary volume *P* and the ground truth binary volume *G*, the Dice coefficient is defined as:

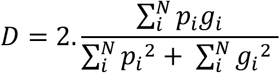

where the sums run over the *N* voxels, of the predicted binary volume *p_i_* ∈ *P* and the ground truth binary volume *g_i_* ∈ *G*. When differentiated with respect to *p_j_* (j-th voxel of the prediction), in order to calculate back-propagated gradients, we get:

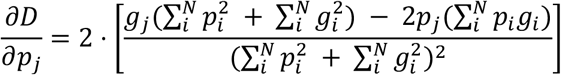

### Model training

All models were trained for 200 epochs. Early stopping was set to 50 epochs where validation loss did not improve to avoid overfitting. The Adam optimizer (Kingma and Ba 2014) was used with an initial learning rate of 5×10^-3^, and a learning rate drop of 0.5 (after 10 epochs where validation loss did not improve). We have selected these training hyperparameters based on experiments.

A multi-contrast network that relies on T1w and FLAIR sequences as inputs were trained. The network was trained on images with various voxel sizes on the whole image, as opposed to patches. Out of the 537 subjects with WMH segmentations used in this study (not including the unseen study used only for testing), 378 (~70%) were used for training, 54 (~10%) for validation during training (from across all the studies), and 105 (~20%) for testing. We tested different optimizers, learning and decay rates. The networks were trained on a V100-SXM2 graphics card with 32G of memory and a Volta architecture (NVIDIA, Santa Clara, CA).

### Segmentation and uncertainty

The MC dropout sampling technique which places a Bernoulli distribution over the network’s weights is implemented to estimate the uncertainty. At test time, by retrieving *N* stochastic outputs, the posterior distribution *p*(*Y*|*X*) can then be approximated (Gal and Ghahramani 2016). With a set *y* = {*y*_1_, *y*_2_., *Y_N_*} from *p*(*Y*|*X*), the final prediction *ŷ* is obtained for *X* by maximum likelihood estimation:

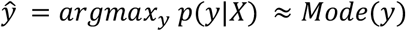

where *Mode*(*y*) is the most frequent element in *y*. This corresponds to the majority voting of multiple predictions.

The uncertainty is estimated by measuring how diverse the predictions are. Both variance and entropy of distribution *p*(*Y*|*X*) can be used to estimate the uncertainty. However, the variance captures the spread among predictions. In this paper we use variance which provides a voxel-wise model uncertainty map:

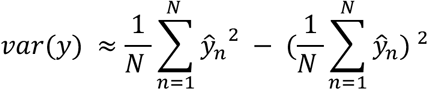

### dWMH and pvWMH segmentation

Distinguishing the dWMH and pvWMH is critical because of their different clinical implications. In this work, we trained a second network to distinguish and segment dWMH and pvWMH. Since multi-class segmentation is more challenging than binary-class segmentation, we trained two separate networks to perform WMH segmentation. The second network was trained on T1w, FLAIR, and binary ground truth as inputs and multi-class ground truth as labels. During inference, the first network generates binary predictions using T1w and FLAIR sequences as inputs. Then, the second network, getting the advantage of the output of the first model, segments dWMH and pvWMH using T1w and FLAIR sequences, and the initial binary WMH segmentation from the first model. The inference pipeline is shown in **Figure 1B**.

### Evaluation on clinical datasets

Our WMH model performance was compared against two other established SOTA segmentation tools: 1) BIANCA from the FSL suite (Griffanti et al. 2016), which is a fully automated and supervised method for WHM detection based on the k-nearest (k-NN) algorithm, and 2) DeepMedic (Kamnitsas et al. 2015, 2017) which is a 3D multi-scale CNN designed with parallel pathways that the second path operates on downsamples images. All segmentation tools used in the analysis were trained and evaluated on the same dataset (training (n = 432) and test (n = 158)) for a fair comparison. To train BIANCA, skull-stripped T1 and FLAIR sequences were used as input data. We used the following optimized BIANCA parameters for training: 1) spatial weighting (sw) = 1, i.e. the data is simply variance normalized; 2) no patch; 3) location of training points = no border location for non-WMH training points, i.e. excluding non-WMH voxels near the lesion’s edge from the training set; and 4) the number of training points = Fixed + unbalanced with 2000 WMH points and 10,000 non-WMH points (Griffanti et al. 2016). Since the output of BIANCA is a probability map of voxels to be classified as WMH, a thresholding step was employed using a 0.9 cutoff to obtain a binary mask.

DeepMedic was also trained using both skull-stripped T1 and FLAIR images. The model architecture included 11-layers (8 layers for the convolutional pathway and 3 layers for final classification), with the convolutional pathway further subdivided into three convolutional pathways. The convolution kernels of the three pathways were the size 3×3×3. The inputs of the three pathways were centred at the same image location, but the second and third segments were extracted from a down-sampled version of the image by a factor of 3 and 5, respectively. DeepMedic was trained with 37×37×37 patches and a batch size of 10 for a total of 700 epochs. The weights of the network were updated by an Adam optimized with an initial learning rate 10^-3^ following the schedule of l0×0.1^epoch^, and L2 penalty weight decay of 10^-4^. A cross-entropy loss is used. Data augmentation was applied during the training procedure through random flipping in the x-, y-, and z-axes with a probability of 50% and the addition of random noise.

### Evaluation metrics

Volume and shape-based metrics were used to evaluate the segmentation performance of the segmentation methods including the Pearson correlation coefficient, the Dice similarity coefficient (DSC), the modified Hausdorff distance (HD95), the absolute volume difference (AVD), recall and F1-score for individual lesions. The Pearson correlation coefficient (Pearson and Galton 1895) was used as a measure of the correlation between the volumes from each segmentation prediction *P*, and volumes from ground truth manually segmented data *G*.

The DSC is a measure of the overlap between two datasets. Given a predicted binary mask, *P* and a binary ground truth volume *G*, the Dice similarity coefficient is defined as:

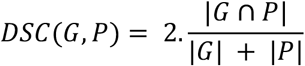

The Hausdorff distance measures how far two surfaces occupying the same space are. Given two sets of points representing objects occupying the same space, *G* and *P*, where *x* ∈ *G* and *y* ∈ *P*, the Hausdorff distance from *P* to *G* is defined as the largest value in a derived set of closest distances between all points.

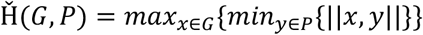

In the above function, ||*x*,*y*|| is the Euclidean distance between points *x* and *y*. Because the Hausdorff distance between the two sets relative to *G* is not equal to the distance relative to *P*, the bidirectional Hausdorff distance is equal to the maximum value between the two directions. A smaller distance is indicative of a greater degree of similarity between the segmentation and the manual tracing. Here we used the 95th percentile instead of the maximum (100th percentile) distance to obtain a more robust distance estimate.

The AVD between the volumes of both ground truth (*G*) and predicted (*P*) images were also computed. An AVD of 0 signifies that the ground truth and the segmentation have the same number of voxels, though it is not indicative of a perfect segmentation. Let *V_G_* and *V_P_* be the volume of lesion regions in *G* and *P* respectively. Then the AVD as a percentage is defined as:

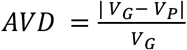

Each individual lesion is defined as a 3D connected component. Given this definition, let *N_G_* be the number of individual lesions delineated in *G*, and *N_TP_* be the number of correctly detected lesions after comparing *P* to *G*. Each individual lesion is defined as a 3D connected component. The Sensitivity for individual lesions (Recall) is defined as:

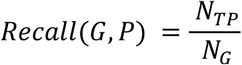

Let *N_FP_* be the number of wrongly detected lesions in *P*. Then the F1-score for individual lesions is defined as:

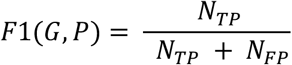

The non-parametric Mann-Whitney U test was employed with an α-level of 0.05 to assess the improvement between our models and all tested methods on these evaluation metrics.

### Clinical adversarial attacks

In order to validate the robustness of our model on data with lower resolution or quality, we generated “clinical adversarial cases” to further test the model’s robustness. These cases included: introduction of noise (to simulate data with lower signal-to-noise ratio ‘SNR’), downsampling of image resolution (to simulate typically short clinical scans performed on low field strength magnets), and different contrasts (to simulate data with different scanners). Noise was introduced using a Rician distribution sampled from two channels of Gaussian noise with a standard deviation of *σ*. Input images were downsampled by a factor of 2 across all dimensions. The image intensities were changed with *γ*. All other SOTA methods were compared to our models on the most challenging adversarial cases, specifically: those with induced noise of *σ* = 0.2, those downsampled by a factor of 2 in each spatial axis, as well as changing image intensity with *γ* = 0.5.

## Results

In this section, we first present our model segmentation results and the accompanying uncertainty maps to highlight the application of the uncertainty for quality control. We then compare the performance of our models to SOTA methods using volume and shape-based evaluation metrics. We also evaluate the second model’s performance across dWMH and pvWMH segmentation. Furthermore, we focus on the evaluation of more challenging mild cases across tested methods. The models are finally tested against generated clinical adversarial cases to evaluate their robustness towards increased noise, lower resolution and different contrasts.

### Uncertainty maps for quality control

**Figure 2** highlights qualitative WMH segmentation results for an example subject based on the Bayesian model with Monte Carlo simulation (*N* = 20 models) to obtain epistemic uncertainty. This example demonstrates instances of both over-segmentation and under-segmentation. It can be observed that the uncertainty map shows high confidence (low uncertainty) in correctly segmented regions and lower confidence (high uncertainty) in mis-segmented regions as highlighted by red (under-segmentation) and green (over-segmentation) arrows. Uncertainty map can thus represent an estimate of segmentation confidence for quality control in both clinical and research settings.

**Figure 2.**
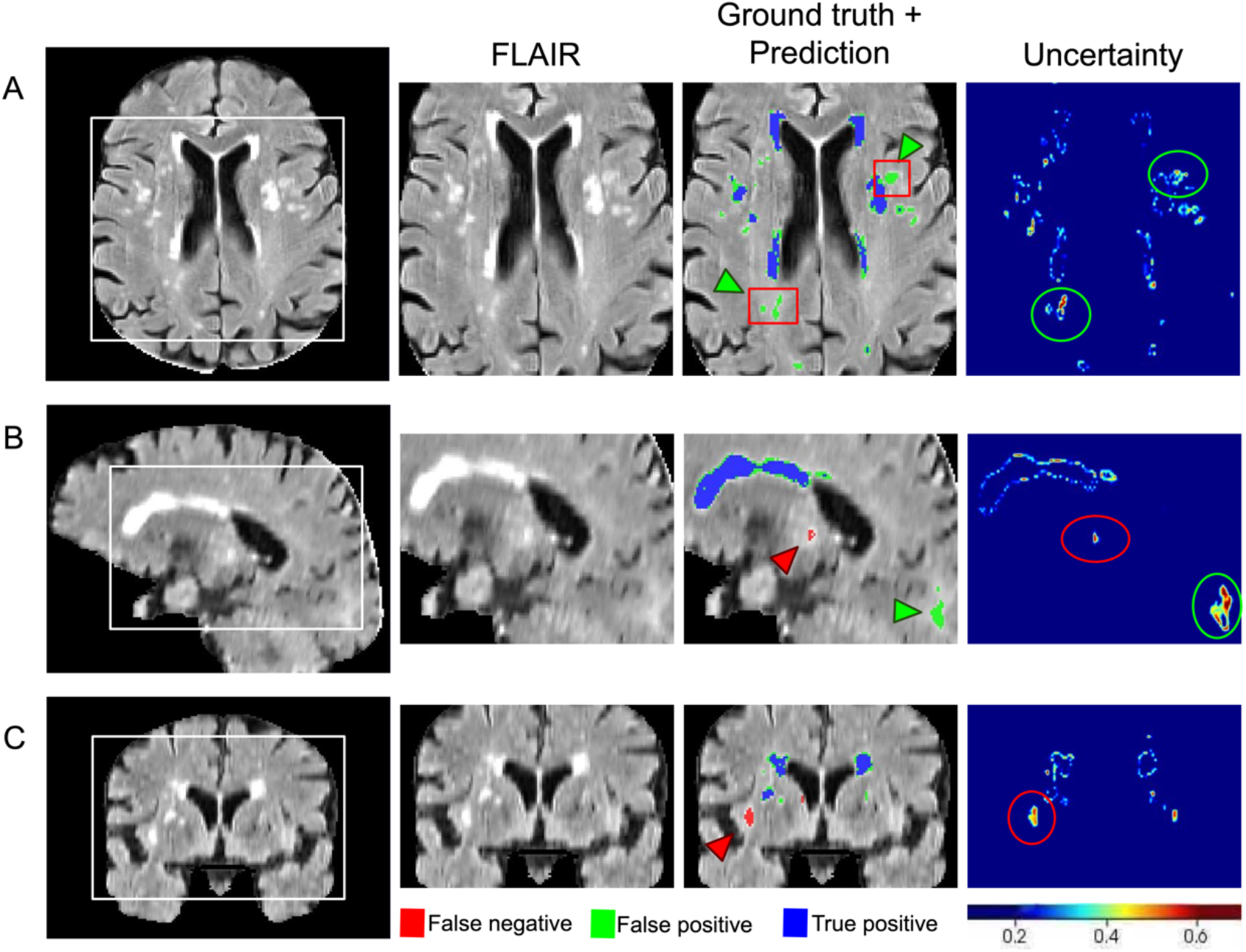
An example of WMH segmentation and uncertainty estimation, showing on a FLAIR scan, the Bayesian model’s prediction and estimated epistemic uncertainty in A) axial B) sagittal, and C) coronal views. Blue represents the overlap between ground truth and prediction, red (and green arrow heads) represent ground truth voxels missing in prediction (under-segmentation), green represent (and green arrow heads) prediction voxels not in the ground truth (over-segmentation). Red boxes represent “false positive” voxels (model predictions) that are indeed positive voxels and were missed in the manual editing of the semi-automated ground truth labels.

### Evaluation of clinical datasets

Average DSC, modified HD95, AVD, recall and F-1 score for HyperMapper Baseline, HyperMapper Bayesian, BIANCA, and DeepMedic for WMH segmentations are summarized in **Table 2** and **Figure 3**. Our Bayesian model had the highest DSC (0.89 ± 0.08) and the lowest HD95 (2.98 ± 4.40 mm) across tested methods. The Bayesian model outperformed the Baseline model, which demonstrates that test-time dropout helps improve segmentation accuracy besides providing an uncertainty map, with the caveat of increased computation time. BIANCA achieved the highest recall, which indicates a high proportion of true positive voxels; however, it produced the lowest F-1 score, which represents low precision or a high number of false positive voxels. Notably, our models had the highest F-1 score which represents lower false positives. DeepMedic and BIANCA had 1.5 to 10 times higher Hausdorff values than our Bayesian model. In addition, our models were faster than BIANCA and DeepMedic. Pearson correlations between manual segmentation volume and volume quantified by the three tested methods are summarized in **Suppl. Figure 1**. Our models and DeepMedic had the highest volume correlations with manual WMH compared to the other tested techniques (*r* = 0.99, *p* < 0.0001), while BIANCA had the lowest volume correlations (*r* = 0.91, *p* < 0.0001). A qualitative comparison between all WMH segmentation methods on one subject is shown in **Figure 4**.

**Table 2.** Evaluation of WMH segmentation on different methods with the following metrics: Dice similarity coefficient (DSC), Hausdorff distance in “mm” unit (modified as 95th percentile) (HD95), absolute volume difference (AVD%), sensitivity (Recall) and F-1 score for individual lesions. ↓ indicates that smaller values represent better performance.

|  | HyperMapper<br>Baseline | HyperMapper<br>Bayesian | BIANCA | DeepMedic |
| --- | --- | --- | --- | --- |
| **Dice similarity coefficient** | 0.892 (± 0.080) | **0.893 (± 0.080)** | 0.604 (± 0.222) | 0.858 (± 0.080) |
| **Hausdorff distance (HD95) (mm) ↓** | 3.045 (± 4.417) | **2.979 (± 4.396)** | 29.684 (± 17.105) | 4.477 (± 6.322) |
| **Absolute volume difference (%) ↓** | **9.819 (± 12.173)** | 9.843 (± 12.134) | 108.970 (± 216.346) | 13.846 (± 14.731) |
| **Sensitivity (Recall)** | 0.762 (± 0.149) | 0.762 (± 0.150) | **0.805 (± 0.168)** | 0.764 (± 0.153) |
| **F-1 score** | 0.752 (± 0.119) | **0.753 (± 0.119)** | 0.199 (± 0.148) | 0.652 (± 0.126) |
| **Time (seconds)** | **10s** | **16s** | 24s | 25s |

**Figure 3.**
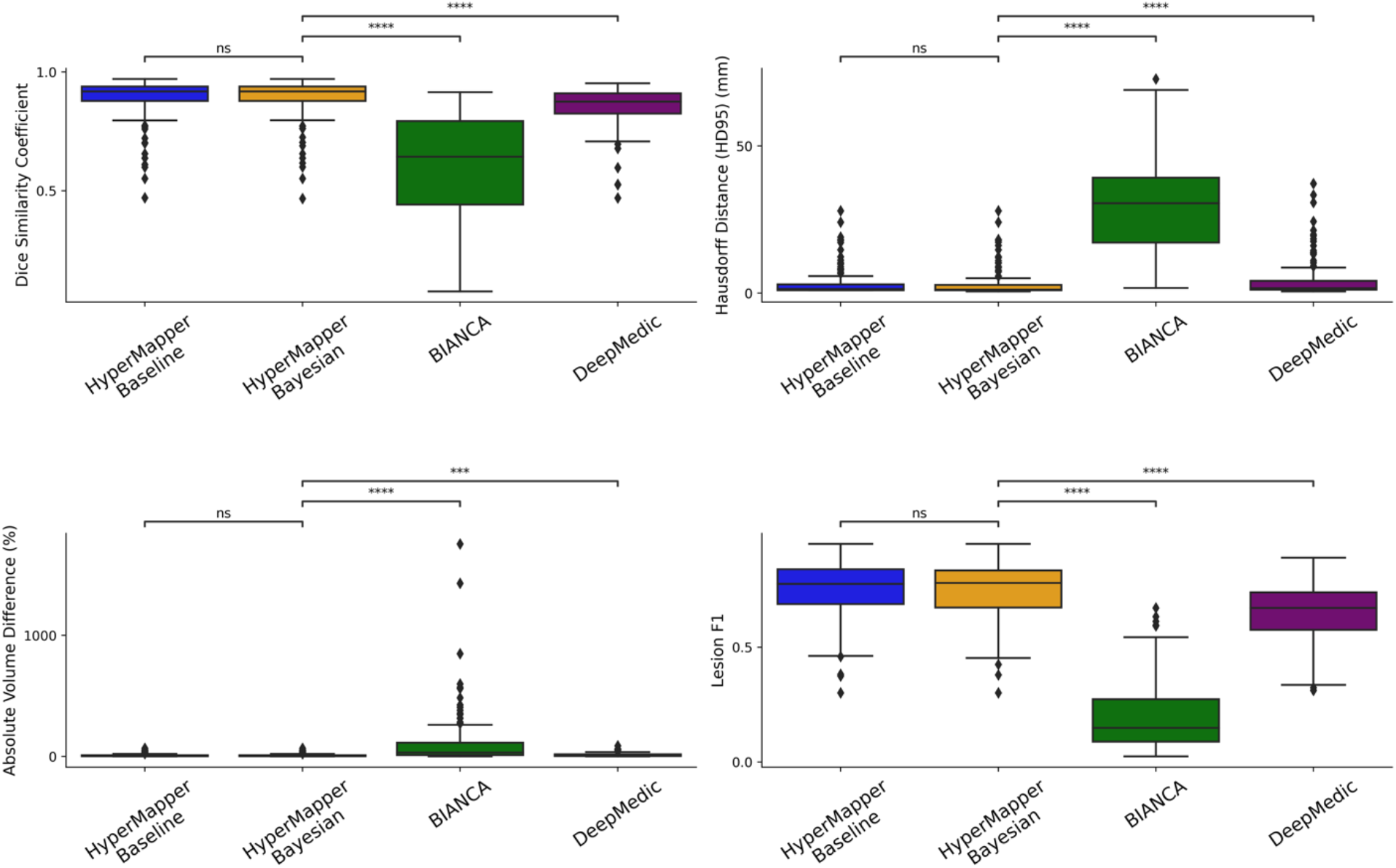
Evaluation of WMH segmentations across tested methods using the following metrics: Dice similarity coefficient, modified Hausdorff distance (HD95), absolute volume difference (%), and Lesion F1. not significant: ns, p < 0.05: *; p < 0.01: **; p < 0.001: ***; p < 0.0001: ****.

**Figure 4.**
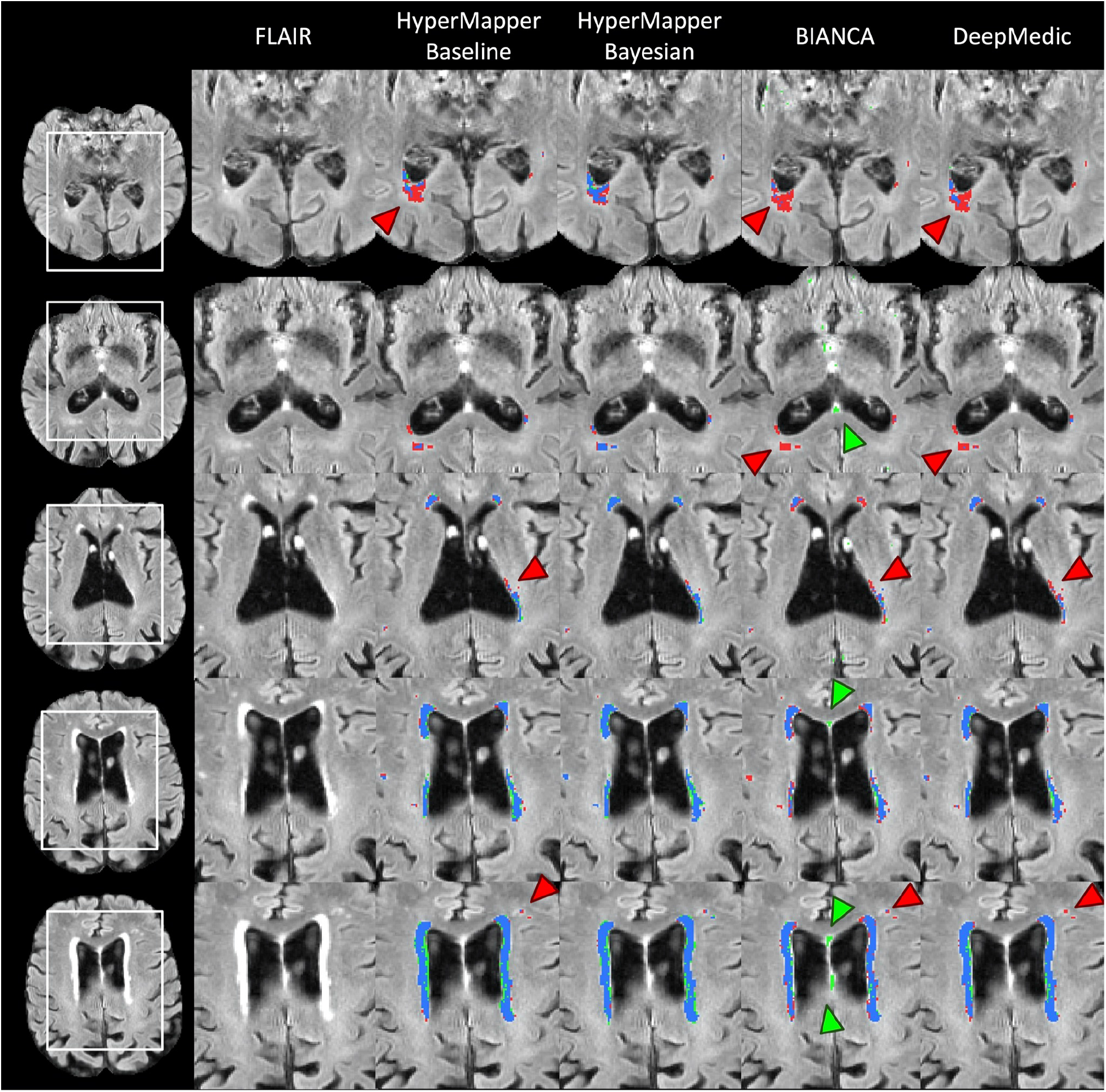
Visual comparison of the tested methods in an example subject. Blue represents the overlap between ground truth and prediction (true positive voxels), red (and red arrows) represents ground truth voxels missing in prediction (false negative voxels), green (and green arrows) represents prediction voxels not in ground truth (false positive voxels).

The cases with the highest and lowest DSC between the manual segmentations and the Bayesian model’s predictions, along with their uncertainties are displayed in **Figure 5** (A and B, respectively). In both cases, the model was able to accurately segment a wide spectrum of white matter disease burden including both very mild and highly severe WMH. Visual inspection of the case with the lowest DSC demonstrates some of the challenges of this segmentation task, such as overlooking hyperintensities in the parietal lobe and ventricle wall. The majority of these challenges were captured in the corresponding uncertainty maps as the high variance between different test-time models.

**Figure 5.**
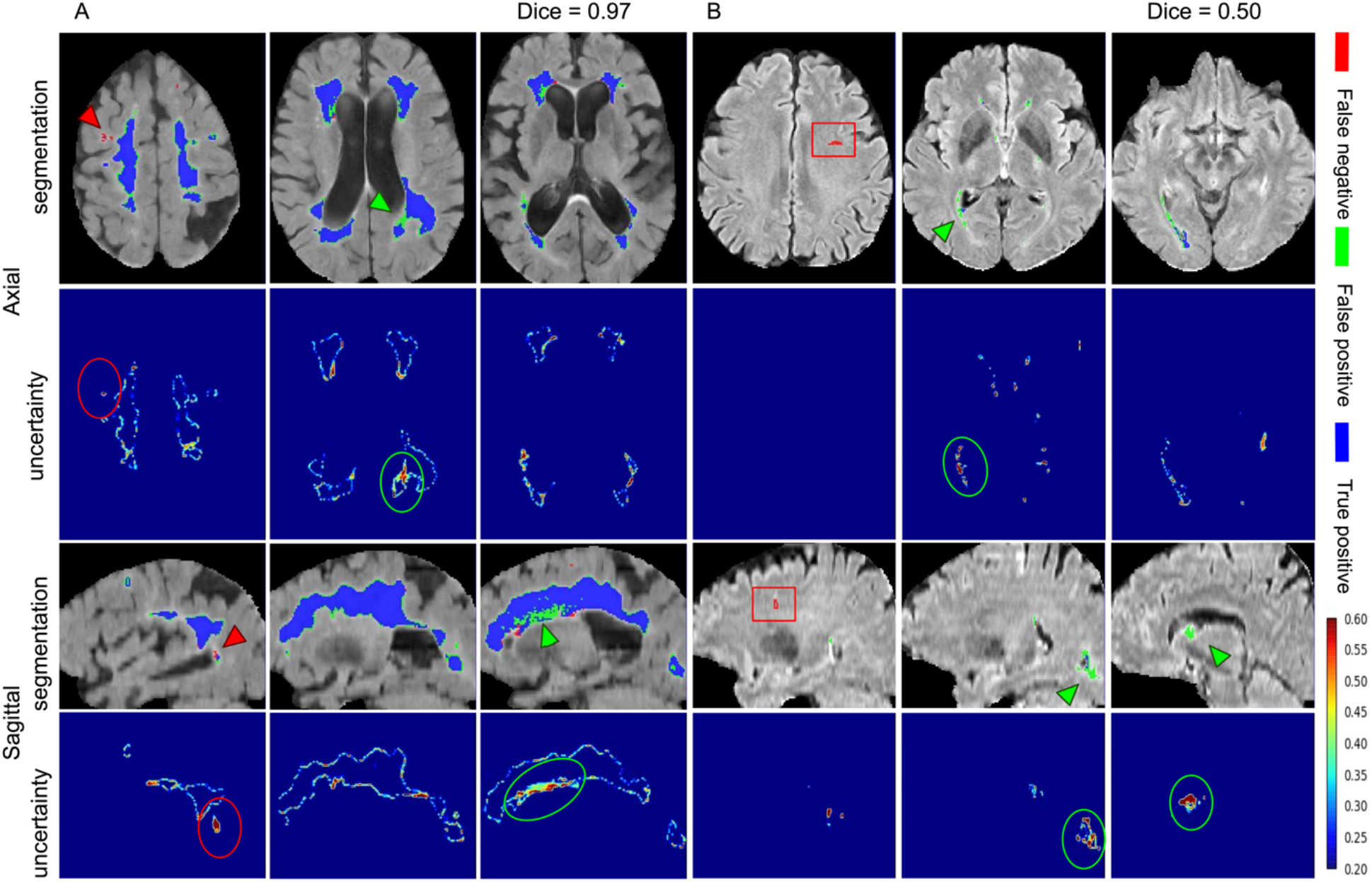
WMH segmentation and uncertainty maps of cases with the highest (A) and lowest (B) Dice similarity coefficients from the test set. Red arrowheads and circles highlight areas of under-segmented and green arrowheads and circles highlight areas that were over-segmented. Red boxes represent an enlarged perivascular space (PVS) in the frontal lobe that was mislabelled as WMH in the ground truth data, but accurately not captured by our model as WMH.

While our training data had only one human rater per image, we also calculated all the evaluation metrics (DSC, modified HD95, AVD, Recall, and the F1 score) on an additional dataset of three human raters with a sample of N=20 to evaluate inter-rater differences. The results are summarized in **Suppl. Table 1** and **Suppl. Figure 2**. Although this additional dataset did not include all the required input sequences to test our model, the raters achieved a Dice similarity that is higher than the one achieved by our model on our test data (0.94 vs 0.89). While this is not a direct comparison, this result is to be expected since these are expert trained raters, the dataset is homogenous (unlike our unseen test data from other studies) and the sample size is smaller than our test data. It should also be noted that our model’s mean Hausdorff distance (HD95) was lower than the inter-rater HD95, probably due to inconsistencies in manual labelling of very small WMH lesions.

### Evaluation on dWMH and pvWMH

We also evaluated our second model’s capability of classifying both dWMH and pvWMH. This distinction is important in the clinical setting because accurate WMH labeling is a critical feature in classifying vascular dementia. Dice similarity coefficients on dWMH and pvWMH were 0.58 ± 0.22 and 0.87 ± 0.09, respectively. These results demonstrate that segmenting dWMH is more challenging due to their size and location. For instance, pvWMH usually appear larger, brighter, and often form confluent lesions with higher contrast, while dWMH usually appears as small punctate lesions. **Figure 6** highlights WMH multi-class segmentation results for an example subject with the first model’s WMH segmentation, dWMH/pvWMH segmentation, and multi-class ground truth.

**Figure 6.**
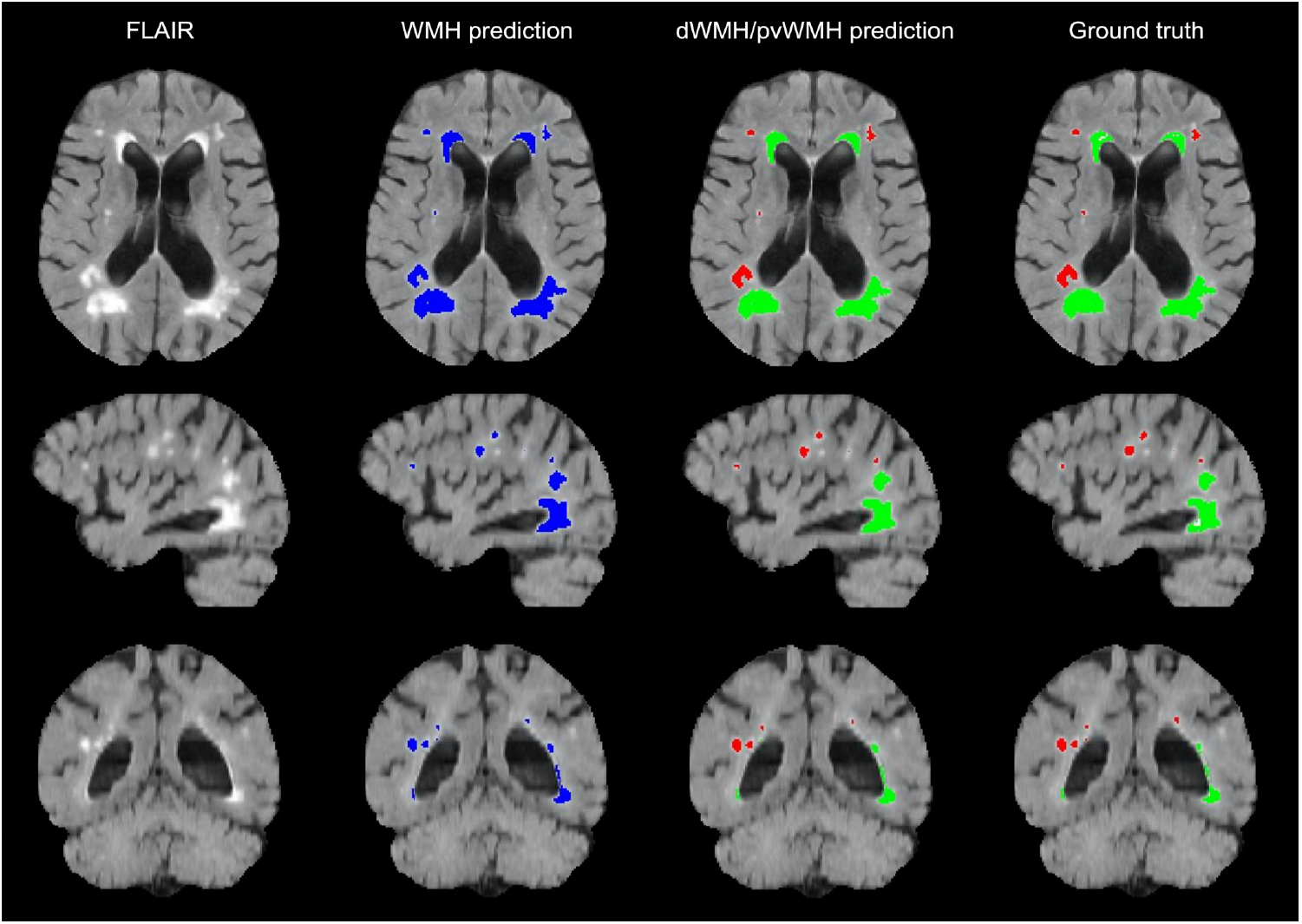
An example of WMH segmentation on a FLAIR scan (axial, sagittal, and coronal view), showing the Bayesian model’s total WMH prediction, dWMH and pvWMH prediction, as well as ground truth labels. Blue labels represent Bayesian model WMH prediction, red labels represent dWMH, and green labels represent pvWMH.

### Evaluation on mild cases

To highlight the clinical and research utility of our models, we further evaluated their performance on cases with mild WMH burden, against other SOTA methods (**Table 3 and Suppl. Figure 3**). **Table 3** shows the evaluation of different methods on 50 cases with mild WMH burden (pvWMH (cc): 1.8 (1.1) and dWMH: 0.27 (0.22)). The results demonstrate that our Bayesian model had the highest performance as assessed by DSC, HD95, and F1-score (0.84 ± 0.10, 5.25 ± 6.51, and 0.721 ± 0.116, respectively). They also highlight the higher complexity of the task and the shortcomings of existing SOTA methods in extracting unique features from much smaller lesions with limited volumes and extent. Therefore, their early detection is clinically important for neurodegenerative populations.

**Table 3.** Evaluation of WMH segmentation on mild WMH cases. ↓ indicates that smaller values represent better performance.

|  | **HyperMapper**<br>**Baseline** | **HyperMapper**<br>**Bayesian** | **BIANCA** | **DeepMedic** |
| --- | --- | --- | --- | --- |
| **Dice similarity**<br>**coefficient** | 0.841 (± 0.097) | **0.842 (± 0.097)** | 0.340 (± 0.067) | 0.779 (± 0.092) |
| **Hausdorff distance**<br>**(HD95) (mm) ↓** | 5.365 (± 6.450) | **5.252 (± 6.505)** | 44.042 (± 9.564) | 8.236 (± 7.190) |
| **Absolute volume**<br>**difference (%) ↓** | **11.298 (± 12.715)** | 11.393 (± 12.812) | 275.970 (± 324.757) | 22.087 (± 20.971) |
| **Sensitivity (Recall)** | 0.712 (± 0.161) | 0.712 (± 0.160) | **0.780 (± 0.187)** | 0.749 (± 0.160) |
| **F-1 score** | 0.720 (± 0.116) | **0.721 (± 0.116)** | 0.096 (± 0.048) | 0.594 (± 0.120) |

### Clinical adversarial cases

Our model’s performance was evaluated on the effects of changes and perturbations in SNR, resolution, and contrast and compared against other SOTA methods (**Table 4**). Cases with the lowest SNR led to a performance drop of our Bayesian model with an average drop of 9.7% in DSC and an average increase of 20% in HD95. While decreases in resolution resulted in an average drop of 9.4% in DSC and an increase of 96% in HD95. Changing image contrast produced a drop of 7.9% in DSC and an increase of 69% in HD95. WMH segmentation results and uncertainty maps for these simulated experiments are shown in **Figure 7** and **Suppl. Figure 9**. To further validate the performance of our model against changes in imaging protocols, we evaluated it against clinical adversarial attacks on a population with mild WMH burden (**Suppl. Table 2**). The results show that the mild WMH cases results in a higher performance drop in our Bayesian model on the DSC (lower SNR: 12.3%, lower resolution: 12.6%, and changing contrast: 10.4%), due to the complexity of small lesions and their higher sensitivity to different attacks such as higher noise and lower resolution.

**Table 4.** Evaluation of WMH segmentation on different adversarial attacks.

|  | Adversarial attacks | HyperMapper Bayesian | BIANCA | DeepMedic |
| --- | --- | --- | --- | --- |
| **Dice similarity coefficient** | Noise (sigma=0.2) | **0.806 ($\pm 0.086$)** | 0.252 ($\pm 0.130$) | 0.639 ($\pm 0.217$) * |
| | downsampled (2x2x2) | **0.809 ($\pm 0.101$)** | 0.348 ($\pm 0.176$) | 0.306 ($\pm 0.208$) ** |
| | Contrast (gamma=0.5) | **0.822 ($\pm 0.102$)** | 0.389 ($\pm 0.184$) | 0.677 ($\pm 0.198$) |
| **Hausdorff distance (HD95) (mm) ↓** | Noise (sigma=0.2) | **3.592 ($\pm 4.686$)** | 31.521 ($\pm 14.999$) | 14.092 ($\pm 13.626$) * |
| | downsampled (2x2x2) | **5.836 ($\pm 6.475$)** | 30.469 ($\pm 18.451$) | 50.049 ($\pm 21.236$) ** |
| | Contrast (gamma=0.5) | **5.047 ($\pm 6.172$)** | 29.488 ($\pm 17.077$) | 10.742 ($\pm 10.5854$) |
| **Absolute volume difference (%) ↓** | Noise (sigma=0.2) | **15.037 ($\pm 13.169$)** | 159.602 ($\pm 278.002$) | 42.627 ($\pm 24.601$) * |
| | downsampled (2x2x2) | **19.745 ($\pm 18.260$)** | 156.712 ($\pm 325.622$) | 77.265 ($\pm 18.012$) ** |
| | Contrast (gamma=0.5) | **24.454 ($\pm 23.178$)** | 176.860 ($\pm 365.183$) | 40.007 ($\pm 24.096$) |
| **Sensitivity (Recall)** | Noise (sigma=0.2) | **0.720 ($\pm 0.136$)** | 0.669 ($\pm 0.168$) | 0.259 ($\pm 0.124$) * |
| | downsampled (2x2x2) | **0.677 ($\pm 0.148$)** | 0.601 ($\pm 0.217$) | 0.072 ($\pm 0.040$) ** |
| | Contrast (gamma=0.5) | 0.601 ($\pm 0.200$) | **0.759 ($\pm 0.186$)** | 0.348 ($\pm 0.191$) |
| **F-1 score** | Noise (sigma=0.2) | **0.505 ($\pm 0.160$)** | 0.139 ($\pm 0.142$) | 0.376 ($\pm 0.134$) * |
| | downsampled (2x2x2) | **0.718 ($\pm 0.010$)** | 0.316 ($\pm 0.169$) | 0.131 ($\pm 0.066$) ** |
| | Contrast (gamma=0.5) | **0.662 ($\pm 0.135$)** | 0.213 ($\pm 0.175$) | 0.469 ($\pm 0.166$) |

\*DeepMedic failed on 1 subject with increased noise, \*\*DeepMedic failed on 62 subjects with lower resolution

**Figure 7.**
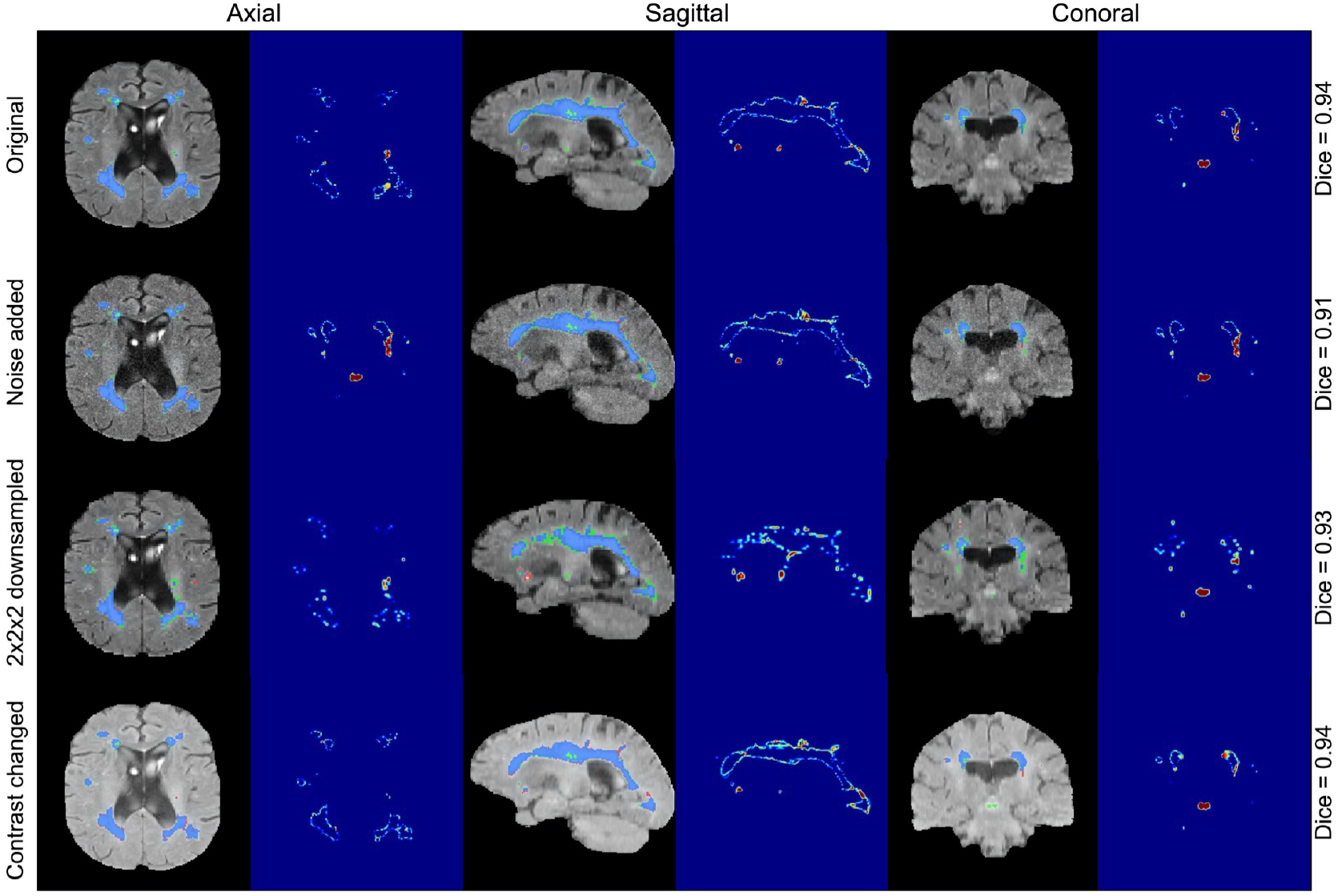
WMH segmentation and uncertainty estimates using our Bayesian model under three types of adversarial attacks applied to the same subject (the addition of noise with a sigma of 0.2, downsampling of resolution by a factor of 2×2×2, and changing contrast with 0.5 gamma).

All tested SOTA methods were compared to our Bayesian model on noisy input data with a sigma 0.2, downsampled data by a factor of 2 across all planes, as well as inputs with contrast changes using a gamma 0.5 (**Figures 8**, **Suppl. Figures 4** and **5**). DeepMedic failed to generate 62 (out of 158) WMH segmentation masks for downsampled adversarial cases, and 1 WMH segmentation for noise induced cases. Failed cases were not included in the analysis or generated figures. We observed that the clinical adversarial cases had significant effects on other SOTA methods as apparent by the substantial decreases in Dice coefficient and increases in Hausdorff distance values (**Table 4**). A qualitative comparison between all WMH segmentation methods on one subject for all three adversarial attacks is included in **Figure 9**. The performance of BIANCA and DeepMedic on the selected subject (**Figure 9**) were high before any attacks (DSC > 0.80) but dropped significantly on adversarial cases.

**Figure 8.**
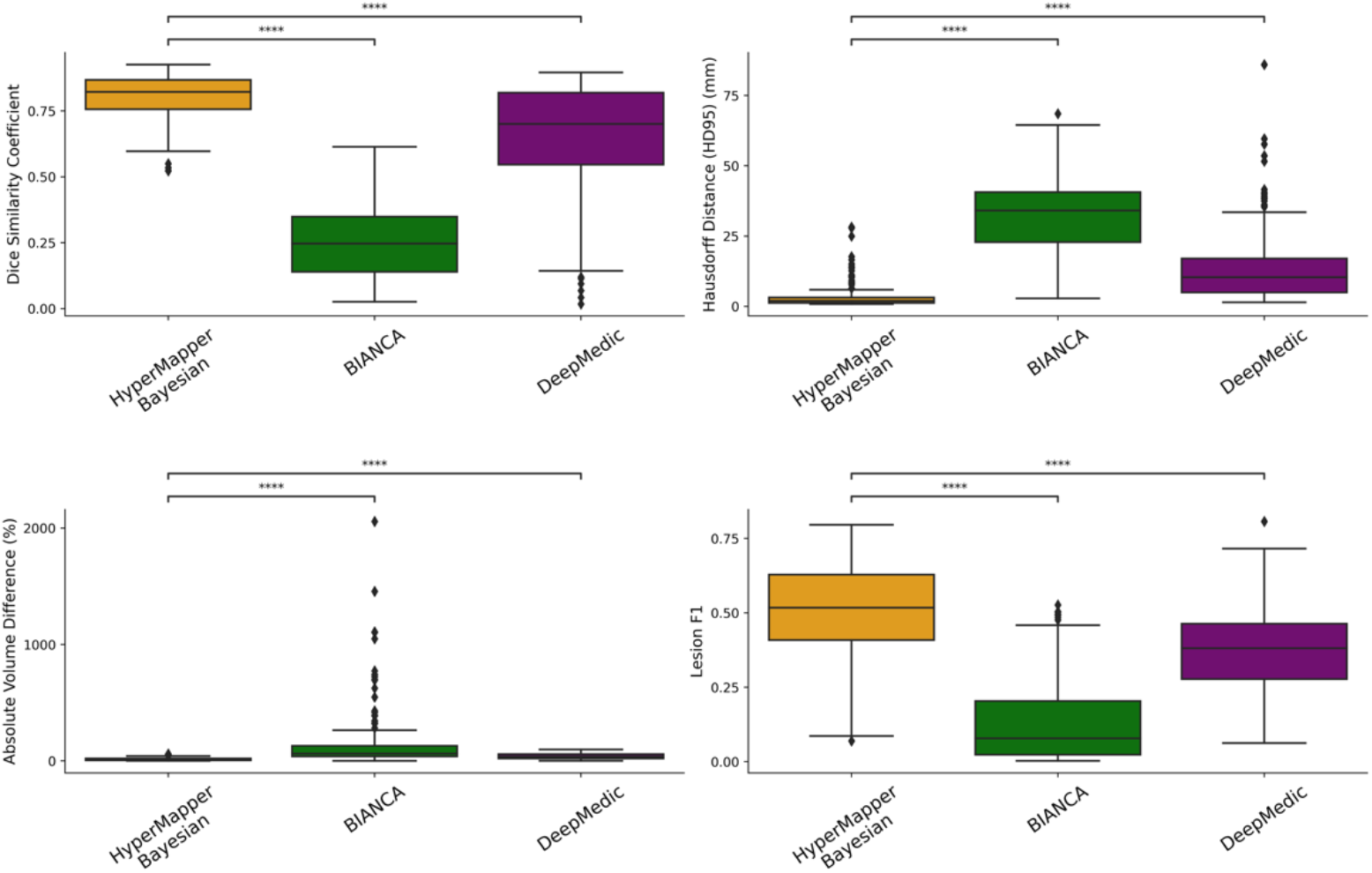
Evaluation of WMH segmentation on cases with increased noise. not significant: ns, p < 0.05: *; p < 0.01: **; p < 0.001: ***; p < 0.0001: ***.

**Figure 9.**
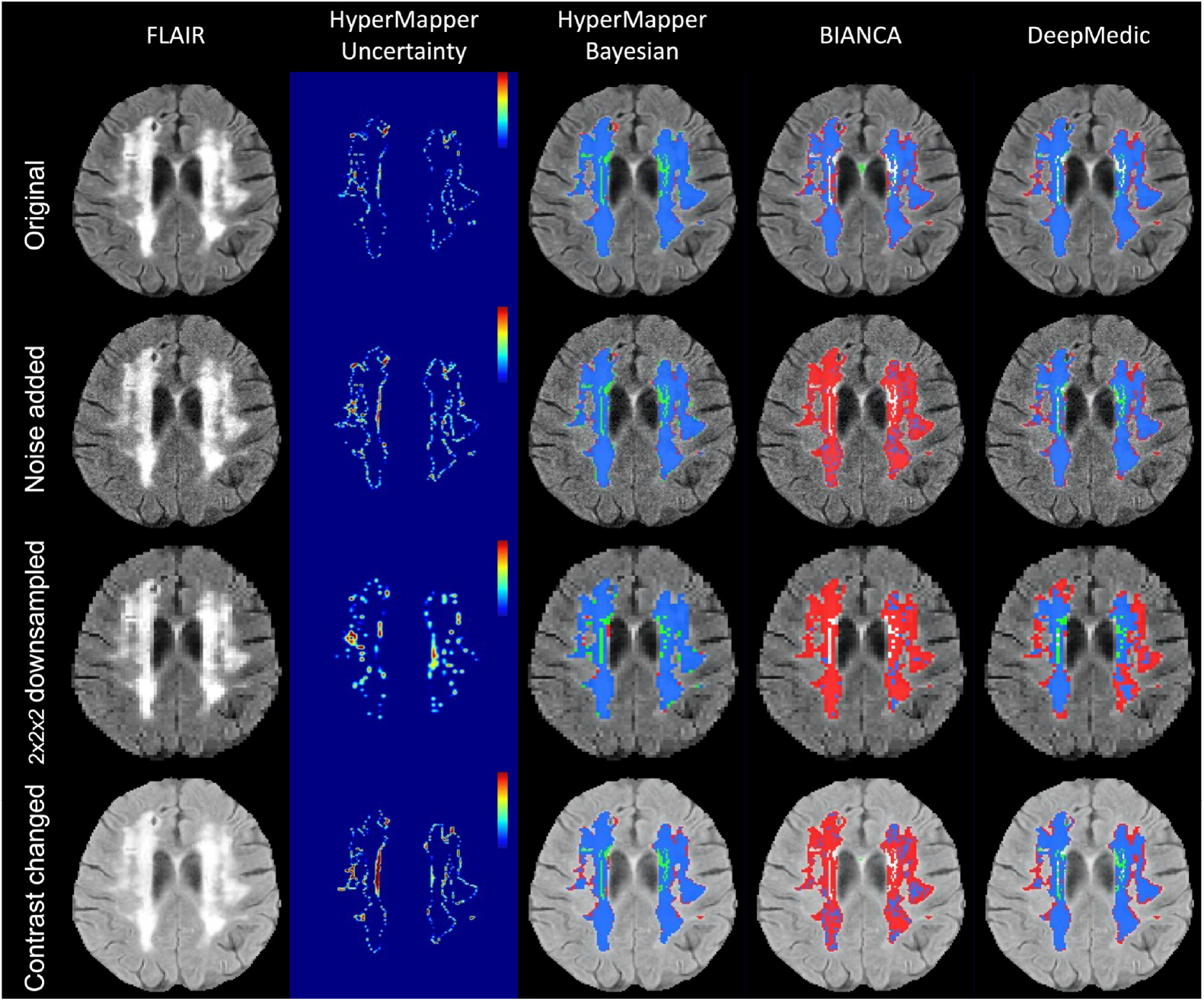
Visual comparison of the segmentation methods under three types of adversarial attacks (the addition of gamma noise with a sigma of 0.2, downsampling of resolution by a factor of 2×2×2, and changing contrast with 0.5 gamma).

Our model had better performances across all metrics for the noise induced cases in comparison to other methods (**Figure 8**). Specifically, the Bayesian model had a significantly higher DSC and lower HD95 relative to other SOTA methods (HyperMapper:-DSC: 0.81 ± 0.09, HD95: 3.59 ± 4.69mm; BIANCA:-DSC: 0.25 ± 0.13, HD95: 30.47 ± 18.45 mm; DeepMedic:-DSC: 0.64 ± 0.22, HD95: 14.09 ± 13.63 mm). Similar to noise induced cases, our Bayesian model had a better performance in comparison to other SOTA methods on adversarial attacks with lower resolution (DSC: 0.81 ± 0.10, HD95: 5.84 ± 6.48 mm). All other SOTA methods encountered a substantial decrease in performance as a result of downsampling the input images. Although DeepMedic’s performance was comparable on in-distribution data, its performance dropped significantly on adversarial cases with lower resolution (DSC: 0.31 ± 0.21, HD95: 50.05 ± 21.24 mm). A similar trend was observed in the contrast permutation adversarial experiments, where a slight drop in accuracy was observed using our Bayesian model (DSC: 0.82 ± 0.10, HD95 5.05 ± 6.17mm); in stark contrast to BIANCA (DSC: 0.39 ± 0.18, HD95: 29.50 ± 17.07 mm) and DeepMedic’s performance (DSC: 0.67 ± 0.19, HD95: 10.74 ± 10.58 mm).

The performance of our models and SOTA methods were further evaluated on adversarial attacks in 50 mild WMH subjects (**Suppl. Table 2, Suppl. Figures 6, 7,** and **8**), demonstrating that our Bayesian model had the highest performance and robustness in comparison to tested SOTA methods across different adversarial attacks. DeepMedic failed to generate 44 (out of 50) WMH segmentation masks for downsampled adversarial cases, and 1 WMH segmentation for noise induced cases. DeepMedic experienced a substantial decrease in performance on adversarial cases with lower SNR (DSC: 0.42 ± 0.20, HD95: 26.64 ± 17.41 mm), lower resolution (DSC: 0.10 ± 0.09, HD95: 76.42 ± 6.63 mm), and contrast changes (DSC: 0.49 ± 0.18, HD95: 20.94 ± 12.35 mm).

## Discussion

This work presented Bayesian 3D CNNs to segment total, deep and periventricular WMH, and estimate model uncertainty to provide a quantitative assessment of confidence in segmentation accuracy. Our models were trained using a heterogeneous multi-site, multi-scanner dataset with different augmentation schemes including the addition of noise, various resolutions, and changes in contrast, to make the model more robust to the challenges that commonly result from different MRI scanners and acquisition protocols. We first highlighted the utility of the resulting uncertainty maps for quality control. Then, we compared our models to available SOTA methods demonstrating our model’s performance on a test dataset including an unseen multi-site study. We further validated the performance of all methods on populations with mild WMH burden. Finally, the performance of our model was validated against SOTA methods on clinical adversarial attacks including lower SNR, lower resolution, and changes in contrast.

### Uncertainty maps for quality control

For estimating model uncertainty, we implemented a Bayesian approach, where MC samples from the posterior distribution were generated by keeping the dropout layers active at test time. Variance over the MC samples was used to provide a voxel-wise model uncertainty map, whereas maximum likelihood estimation over the MC predictions provided the final segmentation. Based on our experiments and visual assessments, the uncertainty maps reliably represented correctly segmented regions with low uncertainty and mis-segmented regions with high uncertainty. Generally, the areas of high uncertainties were localized in the periphery of the segmentations. Furthermore, the uncertainty increased when the model was tested on the out-of-distribution datasets (simulating data with low signal-to-noise ratio, low resolution, and different contrast). Uncertainty is a key concept to highlight scans with lower segmentation accuracy. Uncertainty could therefore be used to either guide the expert or combined with an uncertainty-aware postprocessing method to improve segmentation.

### Evaluation of clinical datasets

While many WMH segmentation algorithms exist due to the importance placed on quantifying WMH in neuroimaging studies and neurodegenerative disorders, these algorithms commonly require manual parameter tuning and are computationally expensive. In addition, many current methods do not produce optimal results in populations with mild vascular lesions. Our models provide an open-source, accurate and robust solution to segment WMH and classifying dWMH and pvWMH that is fast and require no parameter optimization; highlighting their applicability to segment mild WMH burden. Our Bayesian model achieved an average DSC of 0.89 ± 0.08 and HD95 of 2.98 ± 4.40 mm for WMH segmentation in this difficult test dataset.

The Dice similarity coefficient is commonly used to evaluate the overlap between segmentation output and ground truth labels; however, it is less indicative of mismatch when segmenting larger ROIs or outlier voxels. Thus, it should be reviewed in conjunction with surface distance metrics such as the Hausdorff distance as a complementary evaluation metric for measuring boundary mismatches. Although the absolute volume difference (AVD) serves as a measurement of the similarity in voxel counts, it was included as an evaluation metric to provide a fuller context to the segmentation results. To assess over-segmentation and under-segmentation, the values on recall (sensitivity) and F1-score were calculated. Evaluating the Dice similarity coefficient and Hausdorff score, along with AVD, Recall and F1-score, we demonstrated that our proposed model improves on current SOTA methods in terms of accuracy while providing uncertainty maps, performing the task in seconds.

We chose a multi-site and multi-scanner dataset to train our model on ground truth data that were semi-automatically edited by experts rather than automated outputs produced by other algorithms as they are expected to be more accurate. It should be noted that while manual segmentation is often considered the gold standard, it may not necessarily represent the absolute truth. Therefore, some errors in segmentations could be due to inconsistencies in ground truth rather than lower model accuracy. Our proposed method outperformed the semi-automatically edited ground truth in some cases. **Figures 2 and 5** highlighted a few instances where “false positives” are indeed positive voxels and were probably missed in manual editing of the semi-automated lesion segmentation. For instance, **Figure 2A** shows two cases (highlighted by red squares) where errors in segmentations are probably due to inconsistencies in ground truth rather than model false positives, while **Figure 5B** highlights a case where another vascular lesion (enlarged perivascular space ‘PVS’) was mislabelled as WMH in the frontal lobe in the ground truth data (highlighted by red squares), but (accurately) not captured by our model as WMH. It should however be noted that several factors may visually lead to the impression of underestimation of pvWMH by ground truth. For example, partial volume effects at the interface between pvWMH and surrounding normal WM may create a ‘gray’ zone around pvWMH that may be segmented as WMH by CNNs; additionally, increased brightness and lower image contrast may cause lesions to look bigger than their actual size.

When considering the indirect comparison of our model’s performance to inter-rater overlap, while the overlap between raters was marginally higher than the one achieved by the model, this is to be expected in such a complex segmentation task like WMH segmentation, with often small lesions to segment, heterogenous brain anatomy and other confounding vascular lesions. The consistently higher performance of our model on several multi-site studies highlights its robustness against different scan protocols, disease groups, and unseen datasets.

### Evaluation on d/pvWMH and mild cases

Distinguishing the dWMH and pvWMH is important due to different clinical implications such as classifying vascular dementia. While our model achieved a high performance on pvWMH, the task of segmenting dWMH was more challenging due to its shape, size and location. Initial preliminary results showed a worse performance when training a single multi-class model to distinguish dWMH and pvWMH and then generating a binary WMH segmentation (DSC: 0.85). Future work will investigate the potential reasons and how we could improve this single multi-class model performance. Our model’s improvement in performance over SOTA methods was even more apparent in populations with mild WMH burdens. The ability to accurately capture smaller WMH is a more challenging task of lesion detection and quantification; however, it is important for studying and tracking preclinical stages of neurodegenerative diseases and small vessel disease, as it has been shown that WMH at baseline predicts future WMH and is associated with dementia.

### Clinical adversarial cases

Our results on the “clinical adversarial cases” suggest that our Bayesian model is largely robust to cases with varying degrees of noise, downsampling and contrast changes. There are several reasons why our model may be more robust than SOTA methods that we compared against, such as augmentation strategies, residual blocks, dilated convolutions, and skip-connection architecture. Many of the augmentation strategies have been chosen after developing pipelines (using whole-brain images) on multiple applications/tasks to make networks more robust. L-R flipping is one of the most used augmentation strategies. We also performed bias-field correction using the N4 algorithm as a preprocessing step. Tested SOTA methods had widely varying results when segmenting lower SNR data, lower resolution, and data with different contrast than the training set, with substantial volume differences and drops in quantitative metrics or segmentation fidelity. Notably, they failed on the majority of cases with lower resolution, highlighting the sensitivity of some deep learning networks to out-of-distribution data and the need for further validation of the generalizability of these networks. Further improvements of our models may also be necessary in light of some of the weaknesses found, most notably on downsampled and noisy data. For future work, we will investigate incorporating smooth varying maps and other augmentation strategies. In addition to the simulated attacks we employed, there are other challenges that could be tested in future work, such as the simulation of significant subject motion that could have an effect on smaller dWMH.

Though our models performed well compared to other segmentation algorithms, there are still some future improvements that can be made. While our models are trained on a dataset without stroke lesions, the stroke’s penumbra presents with similar contrast to WMH on FLAIR sequences, hence one of the main foci of future work is to train models to distinguish between the two lesions. Future work will also investigate the effect of a combined loss function (for example a weighted cross entropy loss and a weighted dice loss) to handle datasets with varying degrees of contrast and volume sizes.

## Conclusion

We present a robust and efficient WMH segmentation model, which also generates an uncertainty map for quality control. In addition, we present a second model to classify dWMH and pvWMH using the initial total WMH segmentation. We trained our CNN models with expert manually edited segmentations from four large multi-site studies including participants with vascular lesions and atrophy, which represent challenging populations for segmentation techniques, and then tested them on an unseen multi-site study in addition to the four large multi-site datasets. Our segmentation models achieved high accuracy compared to SOTA algorithms on a wide spectrum of WMH burdens, especially mild WMH. Additionally, we used an augmentation scheme to make our model robust to simulated images with SNR, low resolution, and different contrasts. We are making our pipelines and models available to the research community and developed an easy-to-use pipeline with a graphical user interface (GUI) and thorough documentation for making it accessible to users without programming knowledge at: https://hypermapp3r.readthedocs.io.

## Acknowledgements and Funding

This study was funded by the Canadian Institute for Health Research (CIHR) MOP Grant #13129, CIHR Foundation grant #159910, the L.C Campbell Foundation and the SEB Centre for Brain Resilience and Recovery. MG is supported by the Gerald Heffernan foundation and the Donald Stuss Young Investigator innovation award. RHS is supported by a Heart and Stroke Clinician-Scientist Phase II Award. This research was conducted with the support of the Ontario Brain Institute, an independent non-profit corporation, funded partially by the Ontario government. The opinions, results and conclusions are those of the authors and no endorsement by the Ontario Brain Institute is intended or should be inferred. Matching funds were provided by participant hospital and research foundations, including the Baycrest Foundation, Bruyere Research Institute, Centre for Addiction and Mental Health Foundation, London Health Sciences Foundation, McMaster University Faculty of Health Sciences, Ottawa Brain and Mind Research Institute, Queen’s University Faculty of Health Sciences, St. Michael’s Hospital, Sunnybrook Health Sciences Centre Foundation, the Thunder Bay Regional Health Sciences Centre, University Health Network, the University of Ottawa Faculty of Medicine, and the Windsor/Essex County ALS Association. The Temerty Family Foundation provided the major infrastructure matching funds. We are grateful for the support of the Medical Imaging Trial Network of Canada (MITNEC) Grant #NCT02330510, and the following site Principal investigators: Christian Bocti, Michael Borrie, Howard Chertkow, Richard Frayne, Robin Hsiung, Robert Laforce, Jr., Michael D. Noseworthy, Frank S. Prato, Demetrios J. Sahlas, Eric E. Smith, Vesna Sossi, Alex Thiel, Jean-Paul Soucy, and Jean-Claude Tardif. We are also grateful for the support of the Canadian Atherosclerosis Imaging Network (CAIN), and the following investigators: Therese Heinonen, Rob Beanlands, David Spence, Philippe L’Allier, Brian Rutt, Aaron Fenster, Matthias Friedrich, Ben Chow, and Richard Frayne.

## Declarations

### Data Accessibility

The developed algorithm and trained models (network weights) are publicly available at: https://hypermapp3r.readthedocs.io under the GNU General Public License v3.0. An example dataset is included for testing purposes. We have developed an easy-to-use pipeline with a GUI and thorough documentation for making it accessible to users without programming knowledge.

