## supplementary materials for "Deep Bayesian networks for uncertainty estimation and adversarial resistance of white matter hyperintensity segmentation"

### Supplementary Material

**Tables**

**Suppl. Table 1**. Evaluation of WMH inter-rater segmentations (based on three raters on 20 subjects) with the following metrics: Dice similarity coefficient (DSC), Hausdorff distance in "mm" unit (modified as 95th percentile) (HD95), absolute volume difference (AVD%), sensitivity (Recall) and F-1 score for individual lesions. ↓ indicates that smaller values represent better performance.

| | **Dice similarity coefficient** | **Hausdorff distance (HD95) (mm) ↓** | **Absolute volume difference (%) ↓** | **Sensitivity (Recall)** | **F-1 score** |
|---|---|---|---|---|---|
| **Raters** | 0.948 (± 0.072) | 6.866 (± 9.669) | 7.461 (± 10.358) | 0.840 (± 0.158) | 0.853 (± 0.100) |

**Suppl. Table 2.** Evaluation of WMH segmentation on mild WMH cases on different adversarial attacks of different methods with the following metrics: Dice similarity coefficient (DSC), Hausdorff distance in "mm" unit (modified as 95th percentile) (HD95), absolute volume difference (AVD%), sensitivity (Recall) and F-1 score for individual lesions. ↓ indicates that smaller values represent better performance. The failed subjects have been removed for evaluation.

| | **Adversarial attacks** | **HyperMapper Bayesian** | **BIANCA** | **DeepMedic** |
|---|---|---|---|---|
| **Dice similarity coefficient** | Noise (sigma=0.2) | **0.719 (± 0.079)** | 0.132 (± 0.067) | 0.418 (± 0.195) * |
| | downsampled (2x2x2) | **0.716 (± 0.111)** | 0.197 (± 0.106) | 0.100 (± 0.093) ** |
| | Contrast (gamma=0.5) | **0.738 (± 0.099)** | 0.211 (± 0.109) | 0.492 (± 0.178) |
| **Hausdorff distance (H95) (mm) ↓** | Noise (sigma=0.2) | **6.716 (± 6.907)** | 44.124 (± 9.901) | 26.639 (± 17.411) * |
| | downsampled (2x2x2) | **11.500 (± 8.439)** | 47.930 (± 13.844) | 76.415 (± 6.629) ** |
| | Contrast (gamma=0.5) | **9.291 (± 8.640)** | 44.975 (± 12.264) | 20.940 (± 12.346) |
| **Absolute volume difference (%) ↓** | Noise (sigma=0.2) | **19.545 (± 13.938)** | 374.601 (± 417.283) | 66.690 (± 17.729) * |
| | downsampled (2x2x2) | **24.505 (± 24.024)** | 367.883 (± 515.632) | 93.327 (± 6.746) ** |
| | Contrast (gamma=0.5) | **35.918 (± 31.940)** | 442.255 (± 558.893) | 60.716 (± 19.842) |
| **Sensitivity (Recall)** | Noise (sigma=0.2) | 0.672 (± 0.151) | **0.688 (± 0.140)** | 0.248 (± 0.150) * |
| | downsampled (2x2x2) | **0.654 (± 0.142)** | 0.627 (± 0.200) | 0.066 (± 0.002) ** |
| | Contrast (gamma=0.5) | 0.560 (± 0.194) | **0.760 (± 0.153)** | 0.321 (± 0.176) |
| **F-1 score** | Noise (sigma=0.2) | **0.482 (± 0.144)** | 0.042 (± 0.047) | 0.354 (± 0.159) * |
| | downsampled (2x2x2) | **0.707 (± 0.102)** | 0.173 (± 0.094) | 0.123 (± 0.043) ** |
| | Contrast (gamma=0.5) | **0.610 (± 0.119)** | 0.076 (± 0.068) | 0.441 (± 0.147) |

*DeepMedic failed on 1 subject with increased noise, **DeepMedic failed on 44 subjects with lower resolution

### Figures

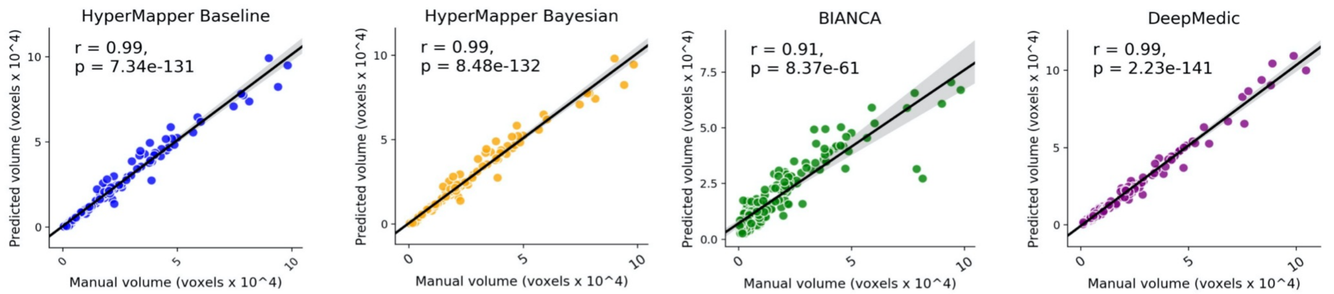


**Suppl. Figure 1.** Pearson R. correlation coefficients and p-values for WMH segmentation between ground truth volumes and predicted volumes using our proposed baseline model (HyperMapper Baseline, blue) and Bayesian model (HyperMapper Bayesian, yellow) as well as two established techniques: BIANCA from the FSL suit (green), and DeepMedic (purple).

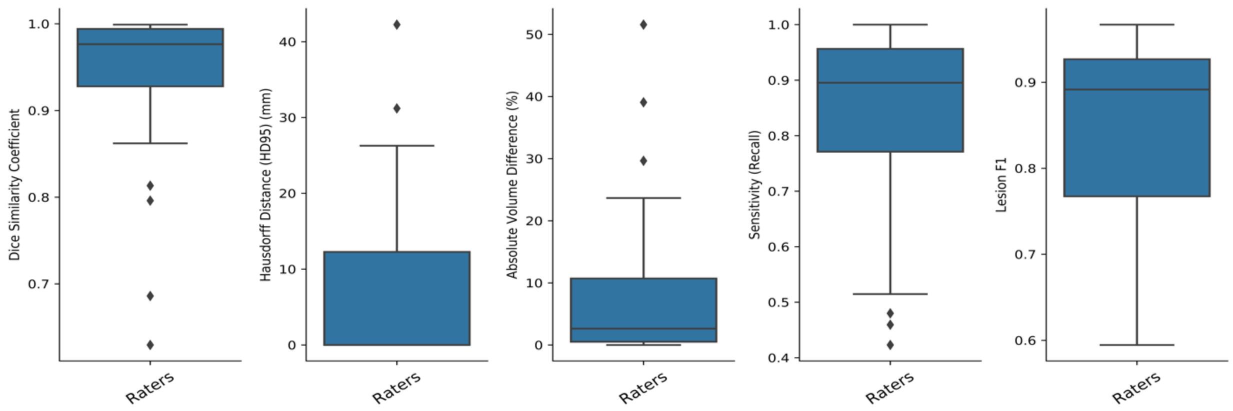


**Suppl. Figure 2.** Evaluation of inter-rater differences on WMH ground truth segmentations (labels), across 20 subjects using the following metrics: Dice similarity coefficient, modified Hausdorff distance (HD95), absolute volume difference (%), sensitivity (recall), and Lesion F1.

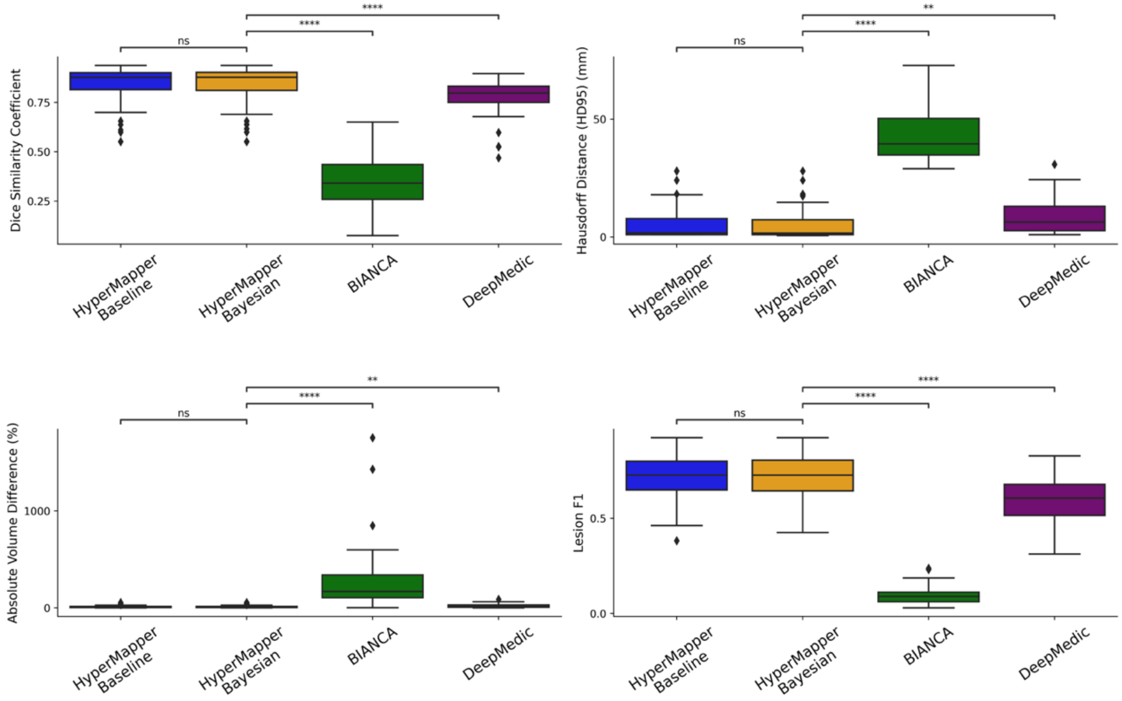


**Suppl. Figure 3.** Evaluation of WMH segmentations on mild WMH cases across tested methods using the following metrics: Dice similarity coefficient, Hausdorff distance (HD95), absolute volume difference (%), and Lesion F1. not significant: ns, p < 0.05: *; p < 0.01: **; p < 0.001: ***; p < 0.0001: ****.

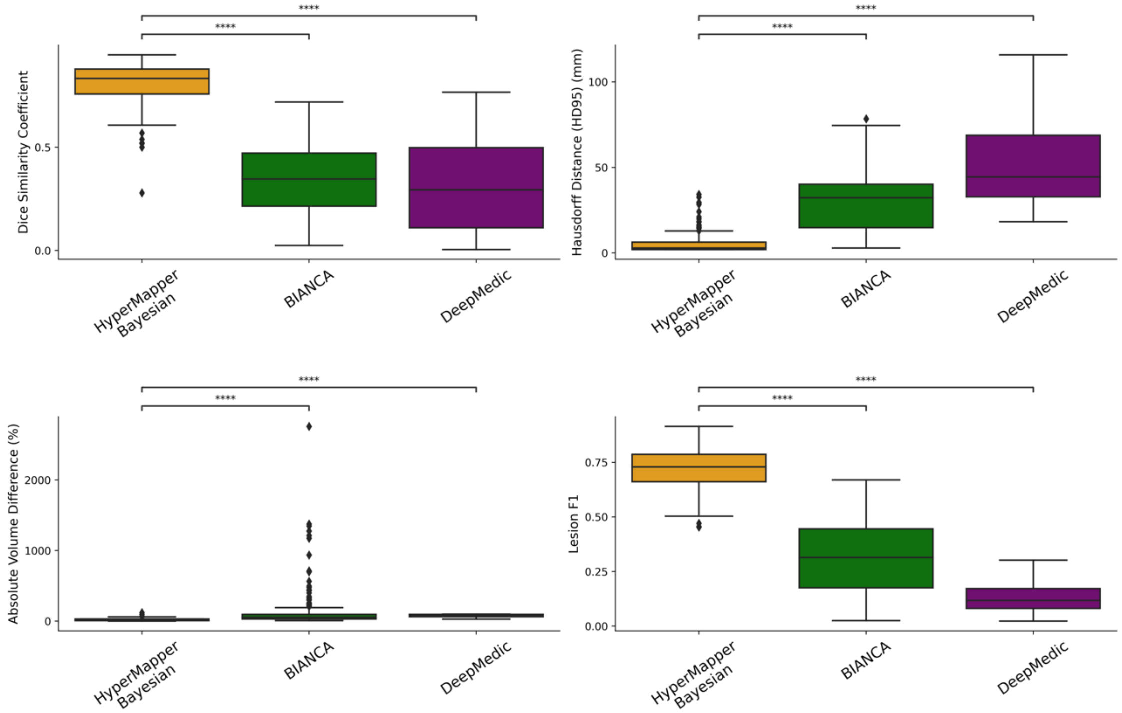


**Suppl. Figure 4.** Evaluation of WMH segmentation on cases with lower resolution. not significant: ns, p < 0.05: *; p < 0.01: **; p < 0.001: ***; p < 0.0001: ***.

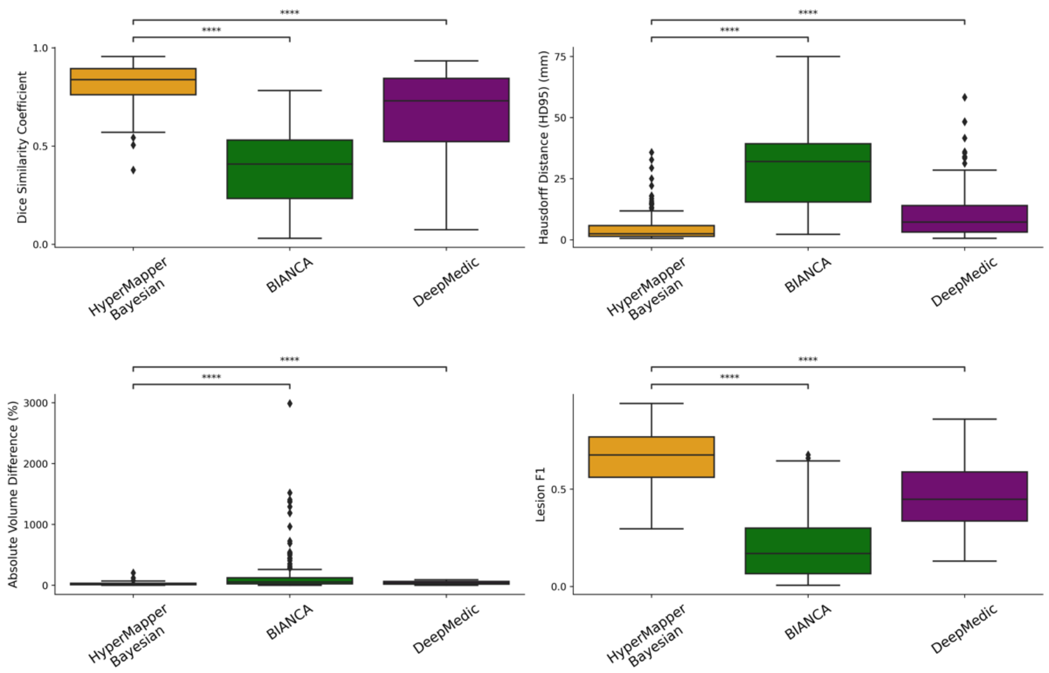


**Suppl. Figure 5.** Evaluation of WMH segmentation on cases with changing contrast. not significant: ns, $p < 0.05$: *; $p < 0.01$: **; $p < 0.001$: ***; $p < 0.0001$: ***.

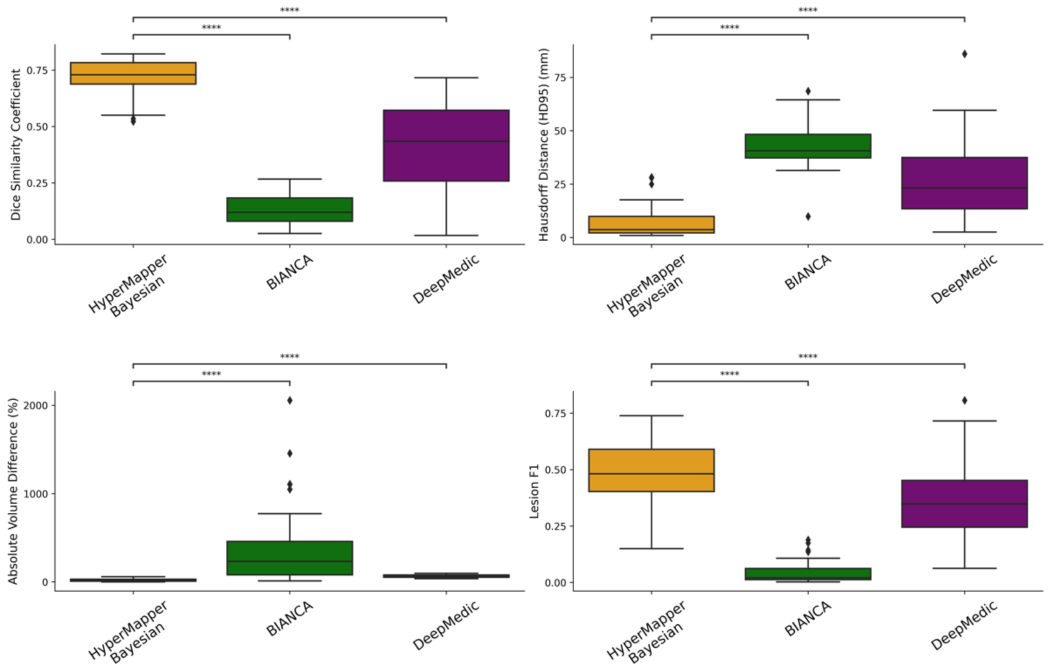


**Suppl. Figure 6.** Evaluation of WMH segmentation on mild WMH cases with increased noise. not significant: ns, $p < 0.05$: *; $p < 0.01$: **; $p < 0.001$: ***; $p < 0.0001$: ***.

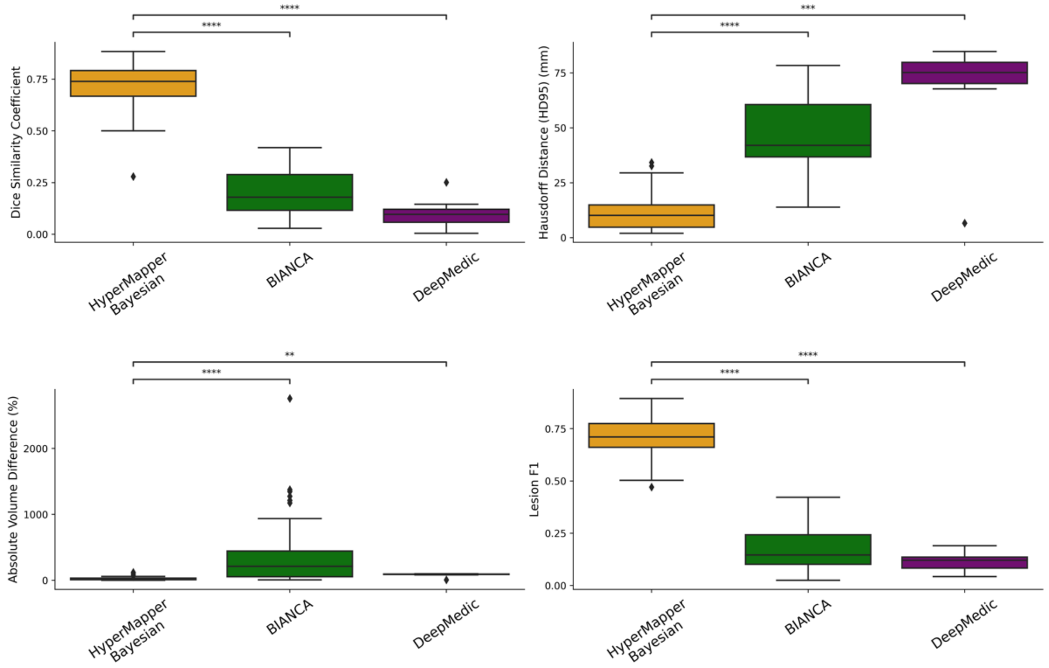


**Suppl. Figure 7.** Evaluation of WMH segmentation on mild WMH cases with lower resolution. not significant: ns, p < 0.05: *; p < 0.01: **; p < 0.001: ***; p < 0.0001: ***.

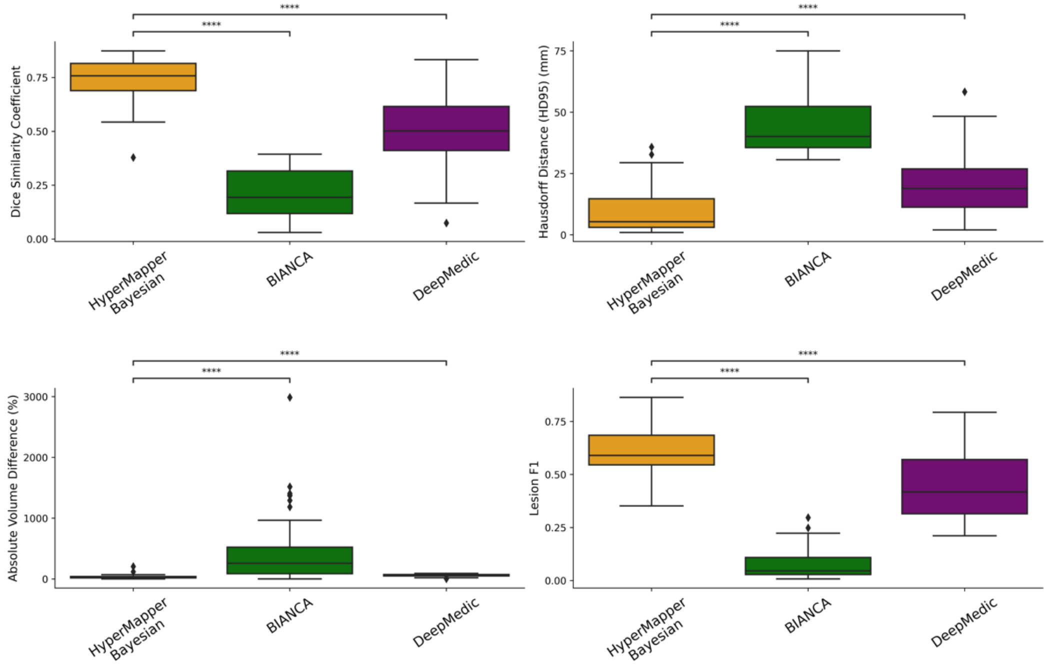


**Suppl. Figure 8.** Evaluation of WMH segmentation on mild WMH cases with changing contrast. not significant: ns, p < 0.05: *; p < 0.01: **; p < 0.001: ***; p < 0.0001: ***.

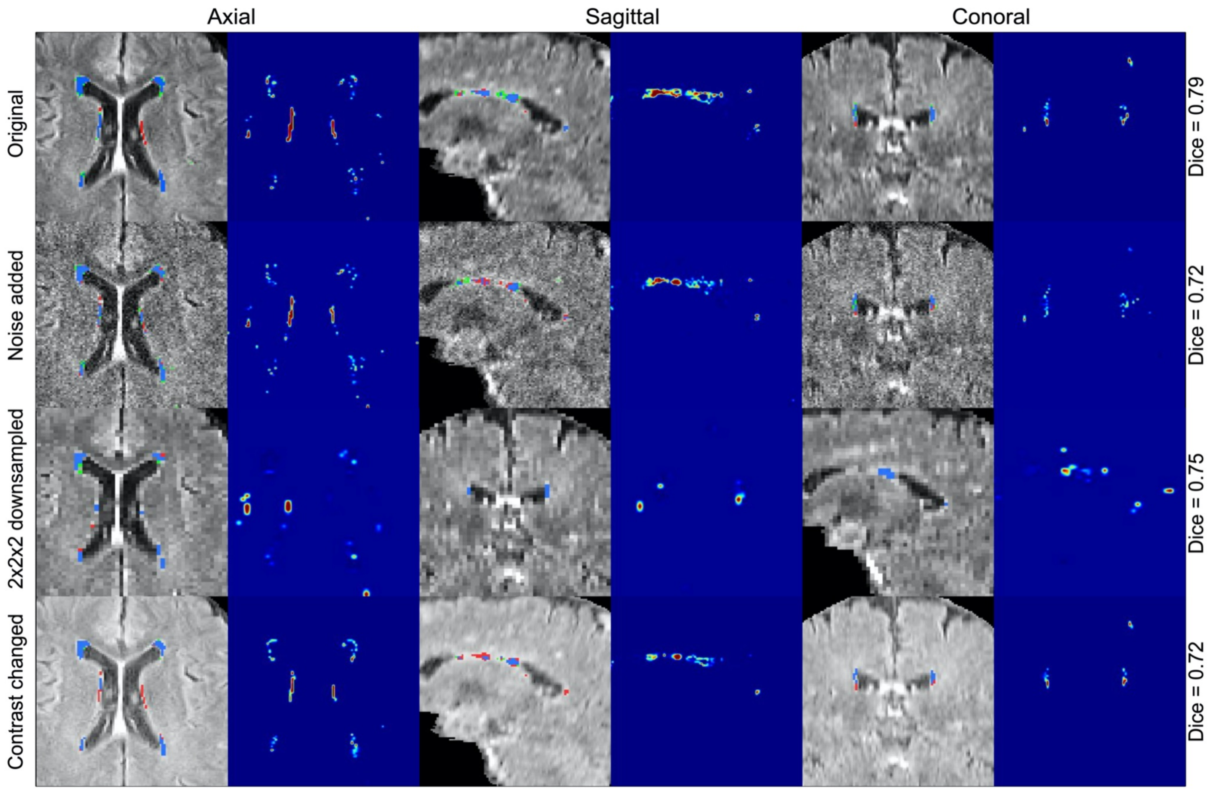


**Suppl. Figure 9.** WMH segmentation and uncertainty estimates using our Bayesian model under three types of adversarial attacks applied to the same subject (the addition of noise with a sigma of 0.2, downsampling of resolution by a factor of 2x2x2, and changing contrast with 0.5 gamma).

**MRI acquisition parameters:**

*Ontario Neurodegenerative Disease Research Initiative (ONDRI) and*
*Medical Imaging Trials NEtwork of Canada (MITNEC) Project C6:*

**3DT1**: TE = (GE: Min full, Philips: 2.3 ms, Siemens: 2.98 ms); TR = (GE: Min, Philips: 9.5 ms, Siemens: 2300 ms); TI = (GE: 400 ms, Philips: 925 ms, Siemens: 900 ms); flip angle = (GE: 11°, Philips: 11°, Siemens 11°); slice thickness = 1 mm; matrix size = 256 x 256; in-plane field of view [FOV] = 256 x 256 mm.

**PD/T2**: TE = (GE: Min full/86, Philips: 10/100 ms, Siemens: 1093 ms); TR = 3000 ms; flip angle = (GE: 125°, Philips: 90°, Siemens 165°); slice thickness = 3 mm; matrix size = (GE: 256 x 256, Philips: 256 x 234, Siemens: 256 x 256); in-plane FOV = 240 x 240 mm; phase FOV= (GE: 75%, Philips: 75%, Siemens: 81%).

**T2 FLAIR**: TE = (GE: 140 ms, Philips: 90 ms, Siemens: 120 ms); TR = 9000; TI = (GE: 2250ms, Philips: 2500ms, Siemens: 2500ms); flip angle = (GE: 125°, Philips: 90° (150° refocus), Siemens 165°); slice thickness = 3 mm; matrix = (GE: 256 x 256, Philips: 256 x 242, Siemens: 256 x 256); field of view [FOV] = 240 x 240.

*Canadian Atherosclerosis Imaging Network (CAIN):*

**3DT1**: TE = 2.3 ms; TR = 9.5 ms; TI = 1400 ms; flip angle = 8°; slice thickness = 1.4 mm; matrix size = 256 x 164.

**PD/T2**: TE = 10.7/102 ms; TR = 2500 ms; flip angle = 90°; slice thickness = 3 mm; matrix size = 256 x 216.

**T2 FLAIR**: TE = 125 ms; TR = 9000 ms; TI = 2800 ms; flip angle = 90°; slice thickness = 3 mm; matrix size = 240 x 217.

*the Language Impairment in Progressive Aphasia (LIPA) and the Vascular Brain Health (VBH):*

**3DT1**: TE = 3.2 ms; TR = 8.1 ms; TI = 650 ms; flip angle = 8°; slice thickness = 1mm; matrix size = 256 x 192.

**PD/T2**: TE = 11.1/90 ms; TR = 2500 ms; flip angle = 90°; slice thickness = 3 mm; matrix size = 256 x 192.
**T2 FLAIR**: TE = 140 ms; TR = 9700 ms; TI = 2200 ms; flip angle = 90°; slice thickness = 3 mm; matrix size = 256 x 192.

**Suppl. Table 3.** Information about MRI scanner and parameters of the dataset.

| Sequence | T1 (SPGR) | | | PD/T2 | | | FLAIR | | |
|---|---|---|---|---|---|---|---|---|---|
| Project | ONDRI/ MITNEC | LIPA/ VBH | CAIN | ONDRI/ MITNEC | LIPA/ VBH | CAIN | ONDRI/ MITNEC | LIPA/ VBH | CAIN |
| In-plane Resolution (mm) | 1 x 1 | 0.9 x 0.9 | 0.94 x 1.1 | 0.94 x 0.94 | 0.9 x 0.9 | 1 x 1.1 | 0.94 x 0.94 | 0.9 x 0.9 | 1 x 1.1 |
| Slice thickness (mm) | 1 | 1 | 1.4 | 3 | 3 | 3 | 3 | 3 | 3 |
| FOV (mm) | 256 | 220 | 240 | 240 | 220 | 230 | 240 | 220 | 240 |
| Field Strength | 3T | 3T | 3T | 3T | 3T | 3T | 3T | 3T | 3T |
